# Ontogeny, species identity and environment dominate microbiome dynamics in wild populations of kissing bugs (Triatominae)

**DOI:** 10.1101/2020.06.05.135871

**Authors:** Joel J. Brown, Sonia M. Rodríguez-Ruano, Anbu Poosakkannu, Giampiero Batani, Justin O. Schmidt, Walter Roachell, Jan Zima, Václav Hypša, Eva Nováková

## Abstract

**Background:** Kissing bugs (Triatominae) are blood-feeding insects best known as the vectors of *Trypanosoma cruzi*, the causative agent of Chagas’ disease. Considering the high epidemiological relevance of these vectors, their biology and bacterial symbiosis remains surprisingly understudied. While previous investigations revealed generally low individual complexity but high among-individual variability of the triatomine microbiomes, any consistent microbiome determinants have not yet been identified across multiple Triatominae species.

**Methods:** To obtain a more comprehensive view of triatomine microbiomes, we investigated the host-microbiome relationship of five *Triatoma* species sampled from white-throated woodrat (*Neotoma albigula*) nests in multiple locations across the USA. We applied optimized 16S rRNA gene metabarcoding with a novel 18S rRNA gene blocking primer to a set of 170 *T. cruzi* negative individuals across all six instars.

**Results:** Triatomine gut microbiome composition is strongly influenced by three principal factors: ontogeny, species identity, and the environment. The microbiomes are characterised by significant loss in bacterial diversity throughout ontogenetic development. First instars possess the highest bacterial diversity while adult microbiomes are routinely dominated by a single taxon. Primarily, the bacterial genus *Dietzia* dominates late-stage nymphs and adults of *T. rubida, T. protracta*, and *T. lecticularia*, but is not present in the phylogenetically more distant *T. gerstaeckeri* and *T. sanguisuga.* Species-specific microbiome composition, particularly pronounced in early instars, is further modulated by locality-specific effects. In addition, pathogenic bacteria of the genus *Bartonella*, acquired from the vertebrate hosts, are an abundant component of *Triatoma* microbiomes.

**Conclusion:** Our study is the first to demonstrate deterministic patterns in microbiome composition among all life stages and multiple *Triatoma* species. We hypothesize that triatomine microbiome assemblages are produced by species- and life stage-dependent uptake of environmental bacteria and multiple indirect transmission strategies that promote bacterial transfer between individuals. Altogether, our study highlights the complexity of Triatominae symbiosis with bacteria and warrant further investigation to understand microbiome function in these important vectors.

## 1. Background

Kissing bugs (Hemiptera: Reduviidae: Triatominae) are hemimetabolous blood-feeding insects predominantly found across the Americas. They are the vectors of Chagas’ disease (CD; caused by the trypanosomatid parasite *Trypanosoma cruzi*) which the majority of ∼150 known species can transmit to a wide range of mammalian hosts, including humans [1, 2]. There are 11 endemic North American species, whose epidemiological relevance has been overlooked compared to their neotropical relatives. However, multiple recent studies have recorded high prevalence of *T. cruzi* in kissing bugs and reservoir mammals, like packrats, racoons, and opossums [3–5], and others have confirmed cases of autochthonous human CD in the USA [6–10].

Thus *T. cruzi* transmission by native US vectors has become a current health concern, emphasising the need for in-depth understanding of triatomine vector biology.

Hematophagous (blood-feeding) organisms are broadly affected by their associated microbial communities (referred to as the ‘microbiome’). Microbiome diversity, composition, and function directly influence various fundamental aspects of host biology, such as immunity, thermal tolerance, and digestion [11–13]. Nutritionally, blood is rich in proteins and salt, lacks vitamins, and its breakdown releases toxic amounts of haem and urea [14, 15]. Many symbiotic bacteria facilitate blood meal digestion and synthesise essential vitamins, making them important mutualists for their hosts [16–22]. Furthermore, the gut bacteria of hematophagous vectors interact with parasites (like *T. cruzi*) occupying the same niche [2, 23, 24]. The microbiome can potentially impede parasite transmission through direct (competition for resources) and indirect (promoting immune response) interactions [25–31]. The most comprehensive background on hematophagous microbiomes has been derived from mosquitos, ticks, and tsetse flies (reviewed in [32]), whereas triatomine-bacteria associations remain neglected.

Since 2015, twelve studies have investigated triatomine microbiomes using next-generation sequencing (NGS) tools. Unfortunately, the data lacks methodological consistency, having been retrieved from a wide variety of DNA templates, including pooled or individual bodies, entire abdomens, the distal part of the abdomen, whole guts, midguts, faeces, and cultured bacterial colonies [33–44], and often further complicated by other variables (e.g. sex, locality, instar, *T. cruzi* infection status). These studies have collectively described a wide range of factors influencing the microbiomes of different species, but seemingly lack consistent patterns [40]. This disparity may stem from inconsistencies between individual sample sets and methodology, but could also reflect Triatominae true biological characteristics. These include lengthy development times through five nymphal instars [38, 42], complex physiology of the alimentary tract [38], and accessory feeding strategies, like hemolymphagy (feeding on arthropod haemolymph), kleptohematophagy (stealing a blood meal from another triatomine), and coprophagy (feeding on faeces) known to be employed by some triatomine species [2, 45–48].

To date, only a single study [41] has sampled microbiomes from multiple wild populations of any triatomine species. Others have mostly targeted South American vectors in domestic environments or laboratory-reared specimens [34, 38, 39, 43, 44], with little consideration for non-urban systems. In this study, we thus focus on wild populations of 5 *Triatoma* species in southern Texas and Arizona. Sampling triatomines within the nests of a favoured host, the white-throated woodrat (*Neotoma albigula*), substantially increases the probability of an identical blood source (a factor known to influence microbiome composition [49–51]) and allows us to collect all five instars and adults. Furthermore, centring this study on *T. cruzi*-negative individuals eliminates another variable known to influence microbiome composition. Our controlled design thus provides the opportunity to: i) evaluate microbiome development across the full ontogenetic spectrum (first to fifth instars plus adults) in natural populations, ii) determine any relationship between triatomine genetic background and microbiome diversity, iii) examine an environmental effect on microbiome composition in species from multiple distinct geographic areas, iv) determine microbiome specificity among *Triatoma* species from the same microhabitat, and v) identify pathogens acquired through feeding on the vertebrate host.

## 2. Methods

### 2.1 Study Sites and Sample set

290 samples were collected from 3 sites in southern Texas in July 2017 (Chaparral Wildlife Management Area, Camp Bullis and Lackland Air Force base in San Antonio) and 3 sites in southern Arizona in July 2018 (Las Cienegas National Conservation Area, University of Arizona Desert Station in Tucson, and San Pedro Riparian National Conservation Area). Sites were accessed with full written permission from the relevant governing bodies (see Acknowledgements). Individuals of *Triatoma rubida, T. lecticularia, T. sanguisuga, T. gerstaeckeri*, and *T. protracta* were collected from the nests of white-throated woodrats (*Neotoma albigula*). We recorded nest coordinates, developmental stage, morphospecies, engorgement, and sex (adults only) for every individual. All samples were preserved in individual vials with 100% ethanol. Additional samples of adult *T. protracta* and *T. rubida* were provided from two houses neighbouring the University of Arizona Desert Station. Since these were adult individuals attracted by black light and not a permanent domestic infestation, we included them in the study.

### 2.2 DNA extraction and basic molecular analyses

The entire abdomen (comprising the whole gut length) of each individual sample was used as a template for DNA isolation with DNeasy Blood and Tissue kits (Qiagen) according to manufacturer instructions. DNA templates were stored at −75 °C prior to molecular analyses, which included host molecular taxonomy, *T. cruzi* infection status, and 16S rRNA gene amplicon library preparation.

To determine *Triatoma* species identity and phylogenetic distance we used the primers 7432F (5’- GGACGWGGWATTTATTATGGATC- 3’) and 7433R (5’- GCWCCAATTCARGTTARTAA- 3’) to amplify a 663 bp fragment of the *cytB* gene [52]. However, the primer pair 7432F and 7433R failed to amplify a PCR product in samples morphologically identified as *T. protracta*, for which some difficulties with *cytB* sequencing have previously been reported [53, 54]. We therefore designed alternative primers, Tpr_F (5’- CCTACTATCCGCGGTTCCTT - 3’) and Tpr_R (5’- GGGATGGATCGGAGAATTGCG - 3’) using three available *T. protracta cytB* sequences and seven sequences of different *Triatoma* species from GenBank. Under the same conditions published by Monteiro et al. [52] the amplification resulted in 380 bp long sequences. The PCR products were cleaned from primers using Exonuclease I and FastAP (Thermo Scientific) enzymes and Sanger sequenced. Phylogenetic background of the *Triatoma spp.* sample set was reconstructed from aligned sequences using Maximum Likelihood with the best fitting model determined by a corrected Akaike information criterion in jModelTest2 [55]. Representative sequences for each species are available in GenBank under the following accession numbers: MT239320- MT239329.

To eliminate infection status as a variable affecting the host microbiome, all samples were screened for the presence of *T. cruzi* in three PCR reactions (as described in Rodríguez-Ruano et al. [42]). In brief, *T. cruzi* presence/absence was confirmed with the universal primer pair TCZ1/TCZ2, targeting any discrete typing unit (DTU) as described by Moser et al. [56]. Additionally, two primer sets (ME/TC1 or TC2) were used to distinguish different discrete typing units of *T. cruzi* [33, 57]. The representative PCR products of all three primer pairs were Sanger sequenced (as described above) and their identity evaluated based on BLASTn searches to confirm the specificity of the screening process. All bands of the expected size from the PCR products were identified as *T. cruzi* DTUs. 57 *T. cruzi* positive samples were subsequently excluded from the analyses. The complete metadata for the samples used in this study are provided in Additional File 1.

### 2.3 Amplicon library preparation

Extracted DNA templates were used for 16S rRNA gene amplification according to Earth Microbiome Project standards (EMP; http://www.earthmicrobiome.org/protocols-and-standards/16s/). Sample multiplexing was based on a double barcoding strategy with EMP-proposed 12 bp Golay barcodes included in the forward primer 515F [58], and additional 5 bp barcodes (designed in our laboratory) within the reverse primer 926R [58, 59]. Barcodes and primers are available in Additional File 1. The resultant amplicons were approximately 500 bp long, including adapters, barcodes, primer sequences, and approximately 400 bp of the 16S rRNA gene V4/V5 hypervariable region.

### 2.4 Novel 18S rRNA gene blocking primer

Since our previous sequencing of insect-associated microbiomes with the 515F/926R [58, 59] primer pair revealed low specificity towards the 16S rRNA gene (resulting in high numbers of host 18S rRNA gene amplicons) we implemented a custom 18S rRNA gene blocking primer. The eight bp at the 5’ end of the blocking primer 926X (5’ GTGCCCTT**CCGTCAATTCCT**-C3 3’) were designed to specifically match the 18S rRNA gene sequence, while the last 12 bp partially overlaps with the 926R (5’ **CCGYCAATTYMT**TTRAGTTT 3’) annealing site. The C3 spacer modification at the 3’ end of 926X was introduced to prevent any prolongation of the blocker. In addition, the blocking primer was used at 10x higher concentration compared to that of the amplification primers [60]. This concentration disparity results in the 926R primers being outcompeted by the blocker 926X during any possible annealing to the 18S rRNA gene, thus increasing the 16S rRNA gene amplification efficiency.

### 2.5 Library controls & amplicon sequencing

In order to confirm the barcoding output and evaluate any effect of our blocking primer on 16S rRNA gene amplification (e.g. possible amplification bias towards some bacterial taxa), the library contained two types of commercially available microbiome mock communities and three microbiome samples of colony reared *Rhodnius prolixus* sequenced in our previous projects. The mock communities comprised three samples of gDNA templates with an equal composition of 10 bacterial species (ATCC® MSA- 1000™) and three samples with staggered composition of the same 10 bacteria (ATCC® MSA-1001™). Altogether seven negative controls (NC) were used to control for the extraction procedure (2 NC), PCR library preparation (2 NC), and well-to-well contamination (3 NC: PCR water template). The PCR amplicons were cleaned using AMPure XP (Beckman Coulter) magnetic beads and equimolarly pooled based on DNA concentration measured with a Synergy H1 (BioTek) spectrophotometer. Since the bead purification did not completely remove the high concentrations of the blocking primer, the final pooled library was purified using Pippin Prep (Sage science) in order to remove all DNA fragments shorter than 300 bp and longer than 1100 bp. The purified library was sequenced in a single run of Illumina MiSeq using v3 chemistry with 2 × 300 bp output (Norwegian High Throughput Sequencing Centre, Department of Medical Genetics, Oslo University Hospital).

### 2.6 Sequence processing

The raw reads were quality checked (FastQC) and trimmed (necessary for reverse reads due to the reduced end-of-read quality) using USEARCH v9.2.64 [61]. The reads were processed according to the following workflow, implementing suitable scripts from USEARCH v9.2.64 [61]. Pair-end reads were demultiplexed and merged. Primers were trimmed and sequences were quality filtered, resulting in a final amplicon length of 369 bp. The dataset was clustered at 100% identity to get a representative set of sequences for *de novo* OTU picking, using the USEARCH global alignment option at 97% identity match [61]. Taxonomy was assigned to the representative sequences using the BLAST algorithm [62] against the SILVA 132 database trimmed for the SSU rRNA gene [63]. Chloroplast sequences, mitochondrial OTUs, and singletons were removed from the final OTU table using QIIME 1.9 [64].

We analysed the final OTU table as three independent datasets, to make sure that our bioinformatic approach did not influence the results. We made the ‘*basic’* dataset (567 OTUs) by filtering extremely low abundant OTUs (as recommended by Bokulich et al. [65]). We generated the *‘decontam’* dataset (5553 OTUs) from the final OTU table by filtering potential contaminants, using the R package ‘decontam’ (V1.5.0) [66] to systematically identify and discard a total of 118 OTUs (complete list in Additional File 1) with a frequency-based approach combined with the post-PCR concentration of each sample. Three of the computationally identified contaminant OTUs (one assigned to the genus *Sphingomonas* and two to *Geobacillus*) were present in all of our negative controls, comprising 223±195 mean total bacterial reads. We generated the *‘ultraclean’* dataset (183 OTUs) from the *‘decontam’* dataset by employing stringent filtering steps to reduce data complexity, based on our previous experience with insect microbiomes. We retained the OTUs that met the following conditions: first, representing more than 1% of reads in any individual sample, and second, being found repeatedly, i.e. in at least two samples across the dataset.

### 2.7 Statistical analyses

We carried out all downstream analyses on the three normalized datasets using rarefaction at 1000 sequences per sample for ‘*basic*’ and ‘*decontam*’, and 500 sequences per sample for *‘ultraclean’*. We used the ‘vegan’ [67] and ‘phyloseq’ [68] packages in R [69] to calculate community quantitative measures (Richness and Shannon index) and ordination analyses (non-metric multidimensional scaling, NMDS; based on Bray-Curtis dissimilarities). We supported the ordination analyses using PERMANOVA tests with beta-dispersion to determine the significance of the tested factors on shaping the microbiome composition.

To test the effect of ontogeny, we analysed the microbiome communities across host developmental stages in *T. rubida, T. protracta, T. lecticularia*, and *T. gerstaeckeri*. To test for differentiation in microbiome communities with a distinct geographic background, we analysed *T. rubida* samples from two different locations in Arizona. We assessed possible species-specific differentiation by comparing individuals from all 5 species, and by comparing *T. gerstaeckeri* and *T. lecticularia* collected from the same nest in southern Texas (thus eliminating the geographic variable). We evaluated the possible effect of host phylogeny on species-specific microbiome patterns. Using Mantel test (implemented in the R package ‘ecodist’ [70]), we tested correlations between microbiome Bray-Curtis dissimilarities and host genetic distance (obtained using Neighbour Joining analysis with Tamura-Nei model for *cytB* alignment with discarded gap and ambiguous positions). The same approach was used to evaluate the effect of geographic distance among sampling sites calculated in QGIS [71].

## 3. Results

### 3.1 Molecular data

The Illumina MiSeq run generated 11,991,455 reads. With negative controls removed, we retrieved a mean average of 13,883 reads per sample and a median average of 9,398 reads. In our positive controls, we retrieved consistent profiles of expected diversity from the commercially produced mock communities (Additional File 1). Two of the staggered mocks lacked one taxon with 0.04% abundance. The composition of equal mocks was consistently biased towards an overrepresentation of *Clostridium* (2.8 times the expected value of 10%), which led to 0.5-8.6% decreases in other taxa. Within the staggered communities, we retrieved most taxa in the expected proportions (from 0.04% to 44.78%). Three of the low abundant taxa (*Clostridium, Lactobacillus* and *Streptococcus*) were overrepresented. The most underrepresented component was *Rhodobacter* (see Additional File 1). All three *Rhodnius prolixus* positive controls showed consistent profiles: *Enterococcus* (mean(SD) = 86(2)%), *Bacillus* (mean(SD) = 10(1)%), and *Arsenophonus* (mean(SD) = 4(1)%), which matched the results of our previous sequencing runs [42].

Most results focus on 170 *T. cruzi* negative samples from the *‘ultraclean’* dataset (see section 2.6; metadata available in Additional File 1). The corresponding results of ordination analyses from the ‘*basic’* and ‘*decontam’* datasets are available in Additional File 2. Phylogenetic clustering based on Maximum Likelihood (Additional File 3) unequivocally determined the 170 samples from ‘*ultraclean*’ to be *T. rubida* (N=81), *T. lecticularia* (N=13), *T. sanguisuga* (N=15), *T. gerstaeckeri* (N=27), and *T. protracta* (N=34).

### 3.2 Microbiome dynamics & host ontogeny

Host ontogeny is a major factor influencing triatomine microbiomes (Fig. 1A-C). *T. rubida* (the most abundant species in our data) shows a pronounced pattern of diversity loss and *Dietzia* accumulation throughout ontogenetic development. *Dietzia* is absent in our *‘ultraclean’* data from the earliest instars, progresses into some L3s, and then clearly increases in L4 nymphs. In most adults it completely dominates the microbiome, comprising 100% of the reads in some individuals (Fig. 1A). The same ontogenetic pattern exists in *T. gerstaeckeri, T. protracta*, and *T. lecticularia* but was not analysed with statistical support due to smaller sample sizes and some instar unavailability (see Additional File 4). Furthermore, there were significant differences in *T. rubida* microbiome diversity between early and late life stages (6 pairwise comparisons retrieved significant differences at the 0.001 confidence interval, Fig. 1B). First instars had the highest Shannon index value (L1 median average = 2.75) and adults had the lowest (L6 median average = 0.01). In a non-metric multidimensional scaling analysis (NMDS), *T. rubida* microbiomes clustered into significantly distinct groups reflecting their ontogenetic development (Fig. 1C; mean stress ≈ 0.16; PERMANOVA R^2^ = 0.288, p ≤ 0.001, with significant Beta-dispersion F = 3.252, p = 0.014 on 999 permutations).

**Figure 1:**
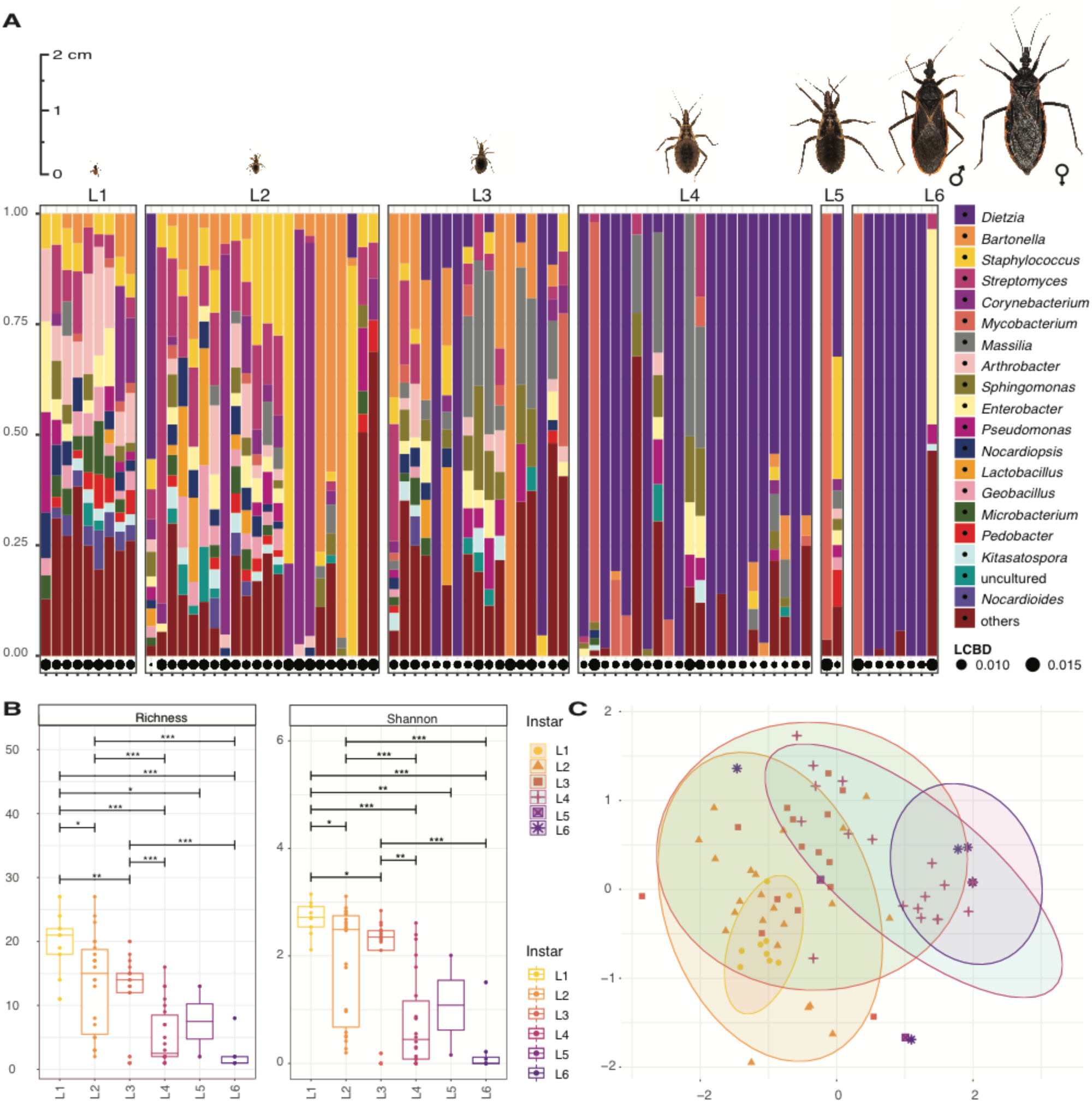
Ontogenetic shift of *Triatoma rubida* microbiomes in sylvatic populations from Arizona. Pictures of each *T. rubida* life stage are to proportional scale. **A:** The microbiome profiles showing the 20 most abundant bacterial genera, listed in decreasing order of abundance to the right of the figure. Each column represents a single individual. LCBD = Local Contributions to Beta Diversity. The size of the circle for each sample indicates the degree of uniqueness for that sample relative to the overall variation in community composition [72]. **B:** Richness and Shannon index (Y axis) for each developmental stage (X axis). Each point represents an individual bug. Length of the horizontal bar indicates the two stages being compared. Asterisks designate significance level as follows *** ≤ 0.001, ** ≤ 0.01, * ≤ 0.05. **C:** NMDS on Bray-Curtis dissimilarity values for microbiomes of different developmental stage. Each colour and shape correspond to a different instar; L6 stands for adults. Each point represents an individual bug. Ellipses are statistically significant at 0.05 confidence interval.

The results presented in Fig. 2 are for two ontogenetic subsets: early (L1-L3) and late (L4-L6) stages, based on their significantly different variance (Additional File 5). There was high among-individual variation in microbiome diversity for all species and instars (Fig. 1A and Additional File 4) but all harboured bacteria from two classes, Actinobacteria and Gammaproteobacteria (Fig. 2). We found species-specific patterns in microbiome composition: only *T. sanguisuga* and *T. gerstaeckeri* contained Acidobacteria and TM6 class bacteria, and only *T. protracta* possessed Bacteroidia in high abundance (particularly the genus *Proteiniphilum*, which dominated some adults; Additional File 4). At the genus level, *Dietzia* dominated the later developmental stages of *T. protracta, T. lecticularia*, and *T. rubida*, yet was completely absent from *T. sanguisuga* and *T. gerstaeckeri* (Fig. 2). Specifically, *Dietzia* comprised 2 OTUs (62% prevalence; abundance median [min-max] = 31.6% [0-100%]) in *T. protracta*, 2 OTUs (77% prevalence; abundance median [min-max] = 79.0% [0-100%]) in *T. lecticularia*, and 3 OTUs (29% prevalence; abundance median [min-max] = 0.0% [0-100%]) in *T. rubida*. In contrast, *T. sanguisuga* was dominated by the genus *Streptomyces* (5 OTUs; 75% prevalence; abundance median [min-max] = 19.1% [0-90%]), as was *T. gerstaeckeri* (6 OTUs; 93% prevalence; abundance median [min-max] = 23.6% [0-90%]).

**Figure 2:**
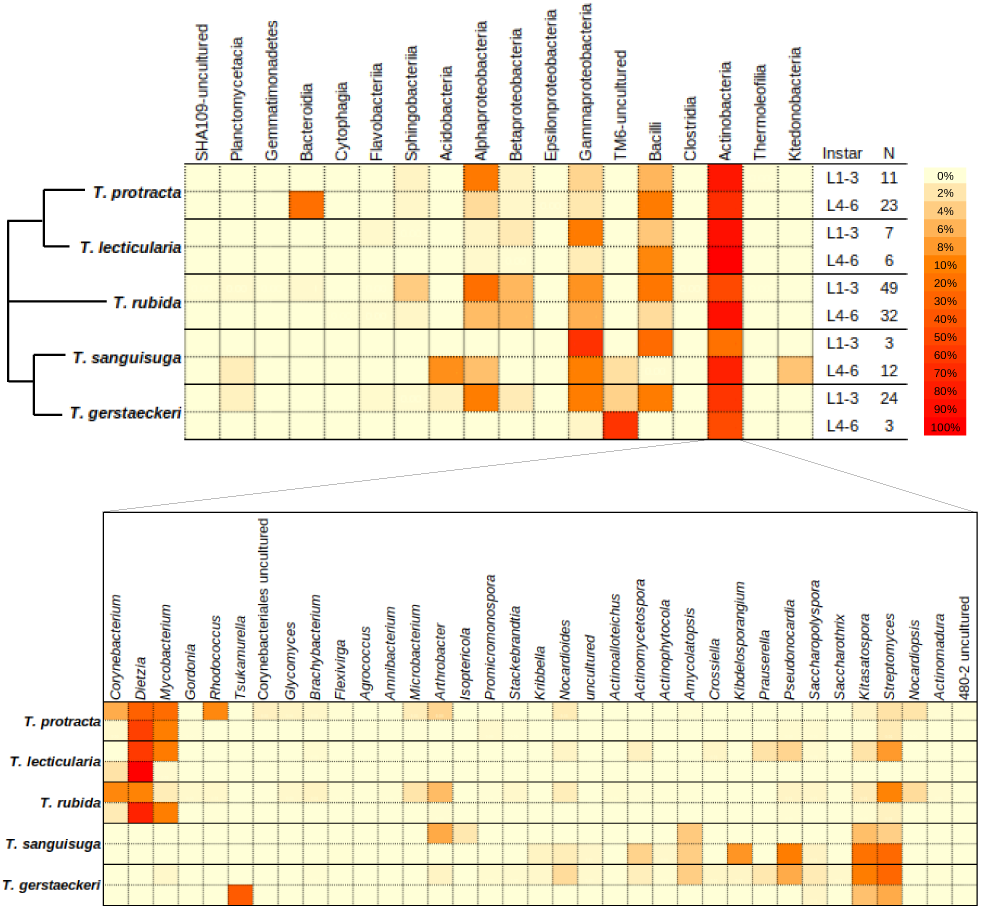
Microbiome profiles of five *Triatoma* species. The top heatmap representing the microbiome profile at order level for early and late instar ranges of all *Triatoma* species. The percentage of each taxon was calculated and normalized by the sum relative abundance per sample, divided by sample size (N). The simplified phylogenetic scheme for the five *Triatoma* species is based on Georgieva et al. [5] and Ibarra-Cerdeña et al. [73], indicating an uncertain position of *T. rubida*. The bottom heatmap represents the genus-level diversity found within the most abundant order (Actinobacteria) present in all species.

### 3.3 Microbiome & host genetic background

We found species-specific differences between the microbiome communities of all five *Triatoma* species (Fig. 3). Evaluation of early instar microbiomes confirmed significant differences among clusters reflecting host species identity (NMDS ordination, mean stress ≈ 0.02; PERMANOVA, R^2^ = 0.18, p ≤ 0.001, Beta-dispersion on 999 permutations: p = 0.034, Fig. 3). To further test host species-specificity in microbiome composition, we specifically compared *T. gerstaeckeri* (17 individuals) and *T. lecticularia* (13 individuals) sampled simultaneously from the same *N. albigula* nest in Chaparral, Texas. These species formed distinct clusters with all individuals included in the analysis (NMDS with mean stress ≈ 0.09; PERMANOVA R^2^ = 0.268, p = 0.002, Beta-dispersion on 999 permutations: p = 0.022; Fig. 4). When we analysed the 23 early instar (L1-L3) individuals, we found the same distinct clusters (NMDS with mean stress ≈ 0.11; PERMANOVA R^2^ = 0.150, p ≤ 0.001, Beta-dispersion on 999 permutations: p = 0.522). They notably differed in microbiome taxonomic composition, with *Dietzia* conspicuously absent from *T. gerstaeckeri* but highly abundant in *T. lecticularia* (Fig. 4).

**Figure 3:**
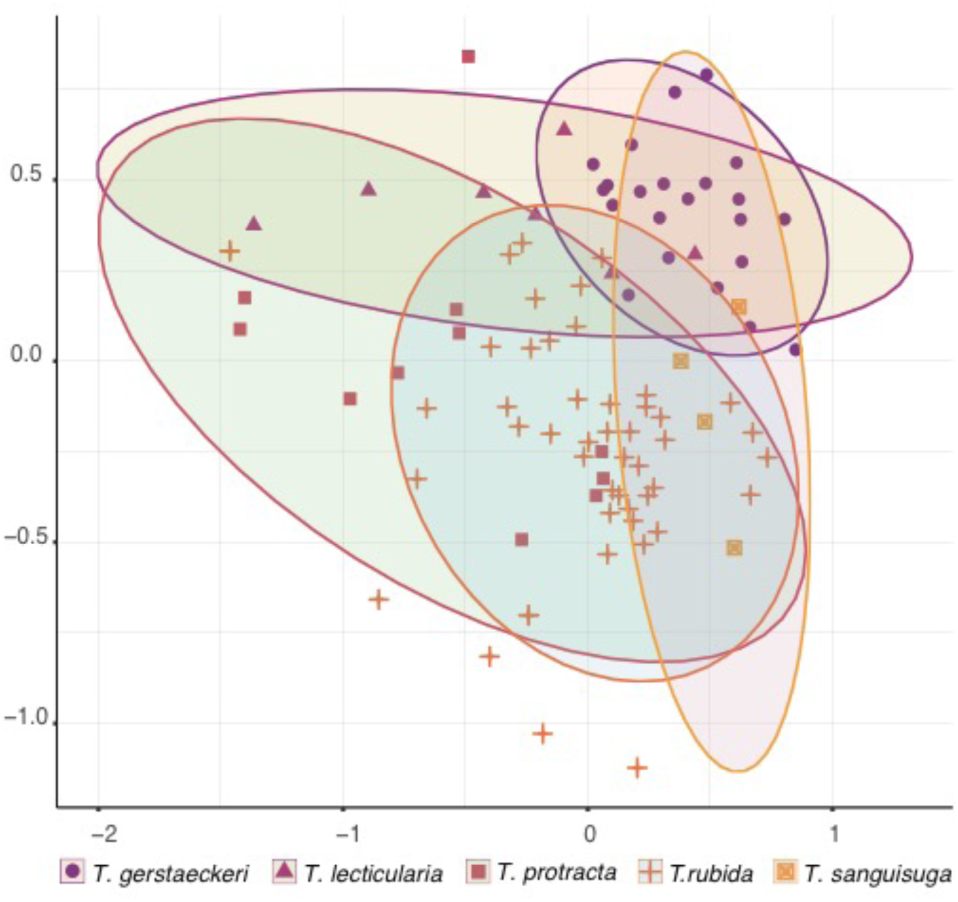
*Triatoma* microbiome species specificity. NMDS ordination for early instars (L1-L3) of all five *Triatoma* species (*T. rubida, T. protracta, T. gerstaeckeri, T. sanguisuga, and T. lecticularia*) based on Bray-Curtis dissimilarities of their microbiome communities. Each point is an individual bug. Each species is represented by a distinct colour and shape shown beneath the x axis.

**Figure 4:**
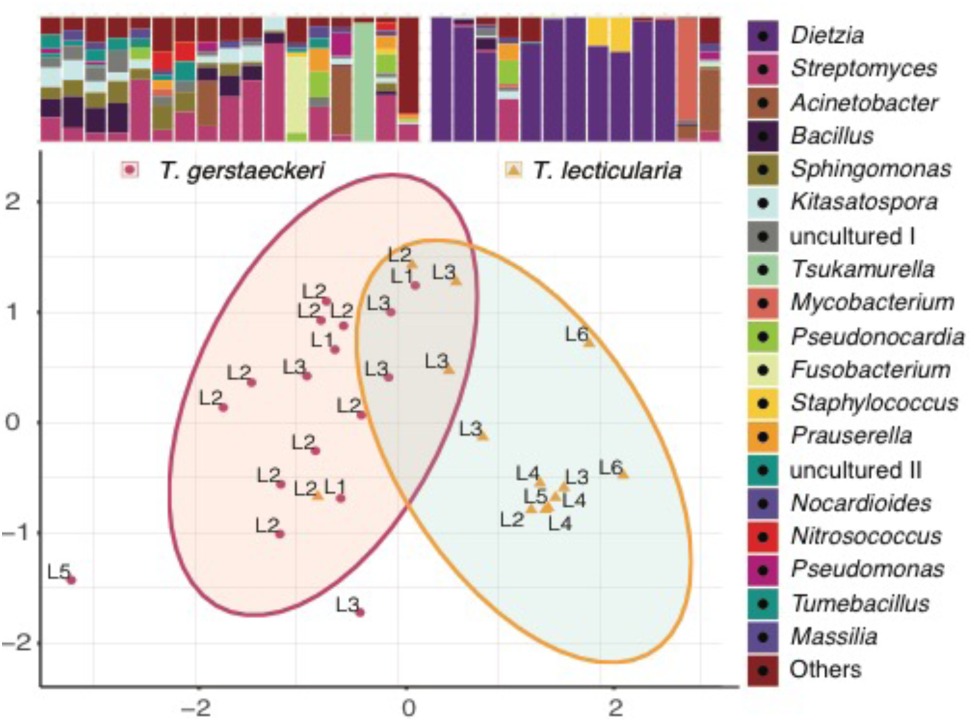
Microbiome species specific differences in a single habitat. NMDS ordination calculated with microbiome Bray-Curtis dissimilarity values from *T. gerstaeckeri* and *T. lecticularia* residing in the same nest in Chaparral, TX. Each point represents an individual bug, with the life stage listed next to it. Statistical ellipses were calculated using a 0.05 confidence interval. Microbiome profiles for both species are provided above the plot, with the 20 top bacterial genera colour-coded and listed in decreasing order of abundance to the right of the plot. “uncultured I” stands for bacteria from the order Bacillales, “uncultured II” for Sphingomonadales.

In addition, we performed two-sided Mantel tests on 94 early instars (L1-L3) to determine if species-specific microbiome differences were a product of host phylogenetic constraint. We identified positive correlations between microbiome dissimilarities and respective host phylogenetic distances (r = 0.29, p ≤ 0.001). We illustrated this result with a Neighbour-Joining tree of Bray-Curtis microbiome dissimilarities that specifically included triatomine species identity and geographic origin (Fig. 5). *T. sanguisuga* and *T. gerstaeckeri* microbiomes were arranged in a single cluster which reflects the host’s close phylogenetic relationship. Microbiomes of *T. lecticularia* predominantly clustered according to host phylogeny (close to *T. protracta* samples), despite the common geographic origin of *T. sanguisuga, T. gerstaeckeri*, and *T. lecticularia. T. protracta* branching was affected by both phylogeny and geographic origin. Overall, both phylogeny and geographic origin partially explain microbiome composition. Analysis of late instars (74 individuals L4-L6) resulted in a notably lower degree of correlation (r = 0.18, p ≤ 0.001).

**Figure 5:**
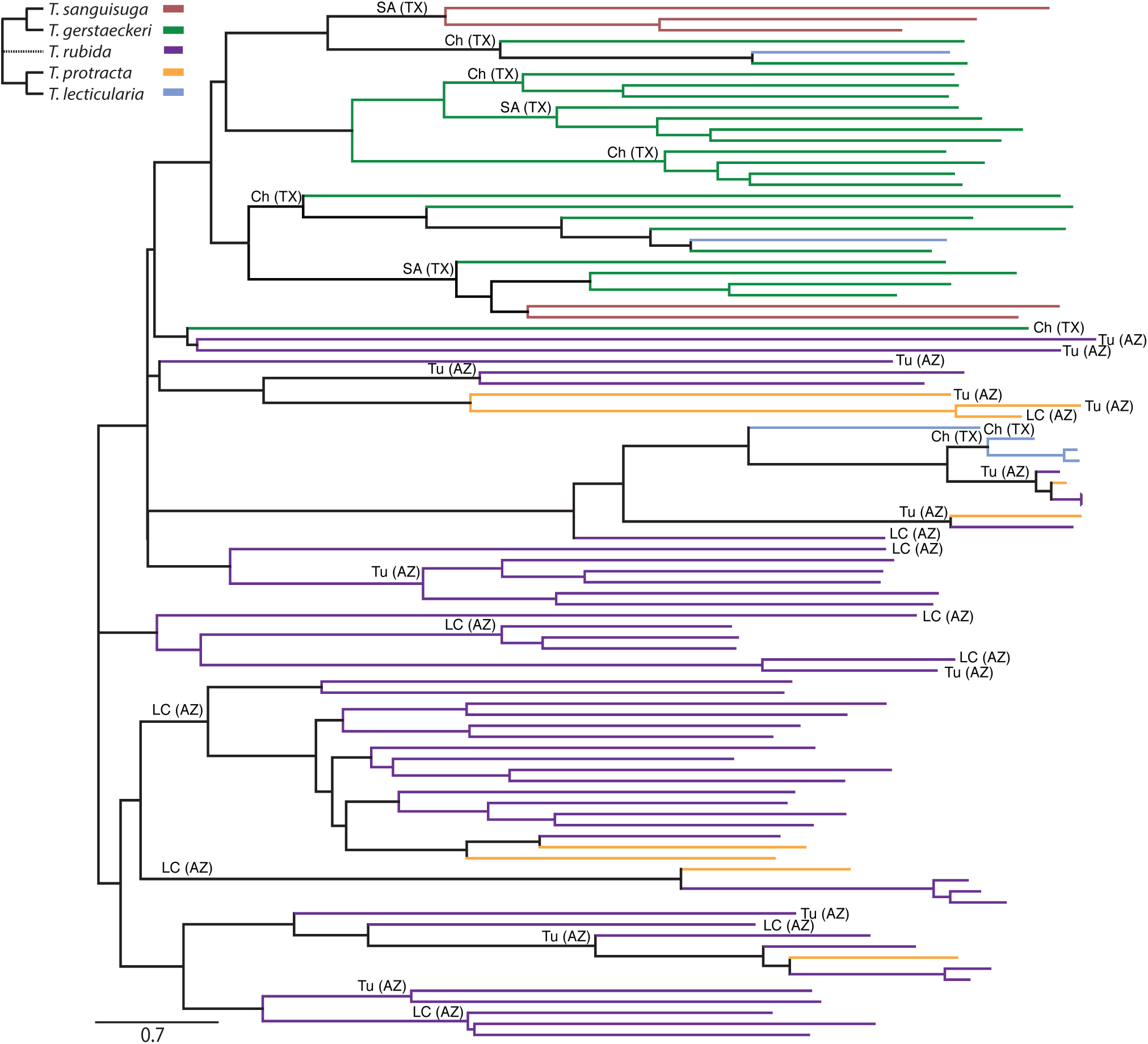
The Neighbour Joining tree of microbiome Bray-Curtis dissimilarities for 94 *Triatoma* early instar individuals, showing microbiome species-specificity is determined by host phylogeny and geographic origin. The nodes are coloured according to host phylogeny. The simplified phylogenetic scheme for the five *Triatoma* species is based on Georgieva et al. [5] and Ibarra-Cerdeña et al. [73]. The dashed line reflects the uncertain position of *T. rubida*. Sample geographic origin is designated as follows: SA (TX) San Antonio, Texas; Ch (TX) Chaparral, Texas; Tu (AZ) University of Arizona Desert Station, Tucson, Arizona; LC (AZ) Las Cienegas National Conservation Area, Arizona.

### 3.4 Microbiome & host geographic origin

Geographic location was a small but significant factor explaining microbiome variation at the intra-species level. We demonstrated this by comparing *T. rubida* from nests in two Arizona locations (Las Cienegas National Conservation Area (LCNCA) and University of Arizona Desert Station (UADS)). We grouped them into early and late instar ranges, to account for ontogenetic changes in microbiome composition, and found their microbiomes significantly differed based on locality. NMDS (mean stress ≈ 0.17) showed statistically significant clusters (PERMANOVA, R^2^ = 0.08, p ≤ 0.001, with non-significant Beta-dispersion on 999 permutations, F = 0.393, p = 0.537; Fig 6).

**Figure 6:**
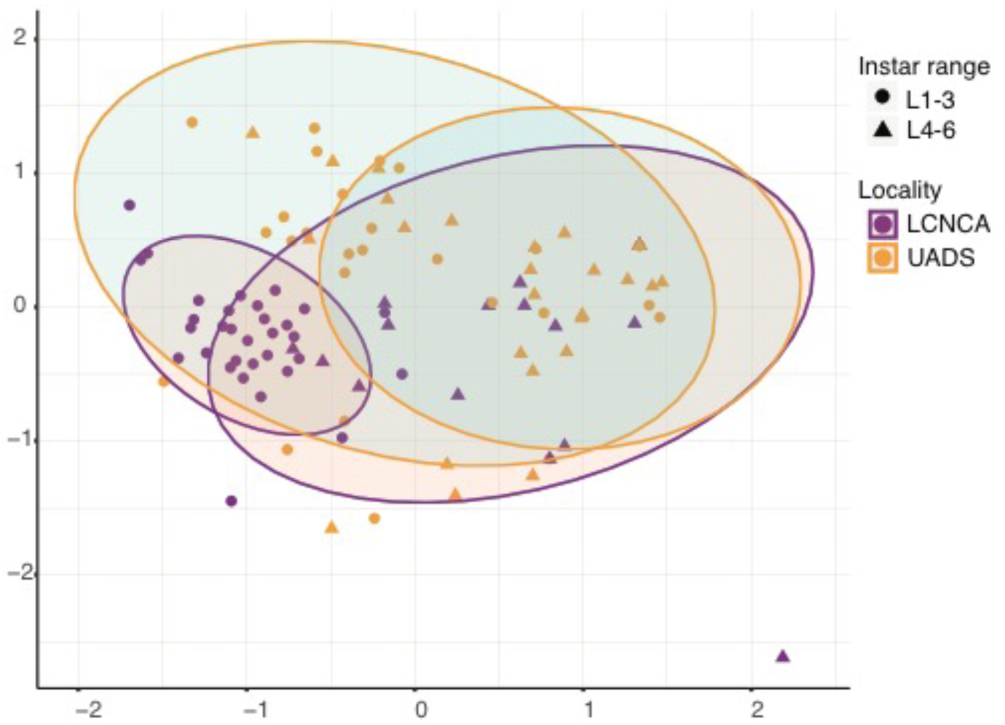
NMDS visualisation of the microbiome differences in *T. rubida* from *N. albigula* nests, sampled in two different locations in southern Arizona. (LCNCA and UADS; represented by different colours). The results are also grouped into early- and late-instar groups (represented by different shapes). The statistical ellipses were assigned using a 0.05 confidence interval.

Furthermore, we analysed the microbiomes of 21 *T. rubida* from 6 different nests within UADS to see if *N. albigula* nests function as natural isolated microhabitats, potentially structuring the population variability amongst *T. rubida* microbiomes. Our results show that microbiome variability reflected the nest origin among early instars (L1-L3), supported with statistically significant clusters in the NMDS analysis (PERMANOVA, R^2^ = 0.45, p ≤ 0.001; Additional File 6) and a modest correlation with geographic distance between the nests (Mantel test, r = 0.164, p = 0.019, at 95% confidence).

### 3.5 Inherited taxa & bacterial pathogens

Sampling across the ontogenetic spectrum allowed us to examine bacterial taxa shared between adult triatomines and their presumed progeny. For this analysis, we only considered adults and early instar individuals originating from the same nest and sharing 100% identity amongst their *coxB* gene sequences. Cross-referencing our ‘*decontam*’ unrarefied data (to deliberately include low abundance taxa that were excluded in the ‘*ultraclean’* dataset) with early and late developmental stages of *T. rubida* (comparison between three L2s and one adult from nest number 4 in LCNCA, AZ) indicated 17 shared OTUs with abundance >0.05% per sample, associated with all individuals. 11 OTUs from *Actinobacteria* represented the genera *Dietzia, Mycobacterium, Corynebacterium, Brachybacterium, Amycolatopsis, Kitasatospora, Nocardiopsis* and *Streptomyces* (4 OTUs), 1 OTU of an uncultured bacterium from *Bacteriodetes*, and 5 OTUs from *Firmicutes* (*Geobacillus, Staphylococcus, Lactobacillus* (2 OTUs) *and Ruminococcus*). For *T. lecticularia* (comparison between three L2 individuals and two adults from nest number 2 in Chaparral, TX), we found 4 shared OTUs with abundance >0.05% per sample associated with all five individuals. These represented the genera *Dietzia* and *Kitasatospora* (*Actinobacteria*), *Bacillus* (*Firmicutes*), and *Enterobacter* (*Proteobacteria*).

Since microbiomes of hematophagous insects often contain acquired bacterial pathogens [74, 75], we examined our data for known pathogenic genera. Two OTUs assigned to the genus *Bartonella* were found in the ‘*basic*’ and ‘*decontam*’ datasets, one of which was retained in the ‘*ultraclean*’ data. This *Bartonella* OTU was highly prevalent (51%) amongst all *T. protracta* and *T. rubida* individuals. Using *Bartonella gltA* gene-specific primers [76], we retrieved 272 bp sequences from *T. rubida* individuals sampled from LCNCA and UADS. We found 99-100% pairwise similarity between the sequences from UADS and *Bartonella vinsonii* isolates from *N. albigula* blood (available in GenBank; KJ719286-7). *Bartonella gltA* sequences retrieved from LCNCA samples were equally similar with *Bartonella vinsonii* isolates from unspecified rodents (AF148491, AF148493, AF148481), suggesting that *T. rubida* acquired pathogenic *Bartonella* from its’ vertebrate hosts (Additional File 7). The representative sequences for *B. vinsonii* found here are available in GenBank under the following accession numbers: MT112947-MT112949.

## 4. Discussion

A wide range of factors have been suggested to determine the microbiomes of triatomines. Species identity, ontogeny, sex, blood meal source, geographical origin, physiological state, and *T. cruzi* infection have all been implicated from the current literature [23, 35, 39, 40]. However, the actual importance of these factors remains inconclusive or controversial. To address this issue within a broader phylogenetic and geographic context allowing for more general conclusions, we designed the first large-scale investigation of triatomine microbiomes, sampling wild populations of five *Triatoma* species. Our data shows that microbiomes of wild triatomines are determined by three main factors: ontogeny, species specificity, and geographic origin, and are also influenced by pathogen uptake from their vertebrate hosts.

### 4.1 Losing diversity: the main ontogenetic shift in microbiome composition

*Triatoma* microbiomes display dramatic compositional shifts from high diversity in first instars towards low diversity dominated by a single bacterium in adults. This general pattern, previously shown only from laboratory reared *Rhodnius prolixus* [42], is well documented in *T. rubida* (our most sampled species) and indicated in three other species (*T. gerstaeckeri, T. protracta*, and *T. lecticularia*). Statistically, ontogeny is the most important explanatory factor of the microbiome dissimilarities found among *T. rubida* individuals. However, there are two caveats to this general pattern of microbiome diversity. Firstly, there is high among-individual variation in richness reduction. For example, some L2s retained highly diverse microbiomes, others showed large reductions in richness reflected by single taxon dominance. The trend towards single taxon dominance increases in later developmental stages (L4 to L6). Among-individual variability was independent of engorgement status (scores are recorded in metadata), suggesting that it is not a product of host physiological state. Furthermore, it suggests that individual triatomines can maintain various different microbiome arrangements (a rich microbiome vs. one dominated by a single taxon). Secondly, *Dietzia* is clearly the dominant bacteria in most late stage individuals of *T. rubida, T. lecticularia* and *T. protracta*, but some of their microbiomes are dominated by other genera (*Mycobacterium, Proteiniphilum*). High single taxon prevalence in late ontogenetic stages likely reflects a real biological process rather than a methodological artefact (e.g. artificial overamplification). We base this assertion on three major points: i) our positive control profiles did not indicate any major preferential amplification in the data, ii) the non-random occurrence of this pattern, i.e. *Dietzia* dominates the late ontogenetic stages of three *Triatoma* species, and iii) concordant results of Mann and colleagues [40] showing 65% of *T. sanguisuga* and *T. gerstaeckeri* adult microbiomes are dominated by a single bacterial taxon, often *Bacillus* or an unspecified *Enterobacteriaceae*.

Currently, we can only hypothesize about the mechanisms underlying the ontogenetic shift from taxon-rich microbiomes in early instars to single taxon-dominated microbiomes in adults. Ontogenetic reduction of microbiome diversity may be random or induced by a specific physiological state, e.g. the moulting process. Insects typically shed the foregut and hindgut linings during moulting, causing loss of symbiotic bacteria in the process [77, 78]. The rich microbial community of first instars (possibly acquired from their eggs [79]) may be periodically shed from the gut with each moulting. After five moulting events, adults thus possess a microbiome with significantly reduced richness. Developmental stage is a key determinant of microbiome composition in other vector species (e.g. ticks [80–82]). However, the relationship between ontogeny and microbiome composition has not been investigated in other hemimetabolous blood-feeding insects (bed bugs or lice; [21]), making generalisable conclusions between biologically similar taxa elusive. Since gut microbiome analysis of natural populations requires killing the specimen, we cannot record microbiome shifts throughout ontogenetic development of a single individual, and thus cannot currently determine whether diversity loss is a permanent change to triatomine microbiomes during their development.

### 4.2 Origin of the microbiome bacteria: inheritance vs environment

The results shown in Figures 2-6 demonstrate that triatomine microbiome composition (especially in early instars) is shaped by host species and locality. For both variables, there was partial overlap in microbiome composition, likely due to the notable degree of among-individual variability (discussed above). Species-specific differences were present even when multiple triatomine species were sampled in the same nest, showing that they are not caused by different environmental sources of bacteria. In theory, they could be explained by multiple different mechanisms, like specific maternally inherited bacteria or differential uptake and retention of environmental bacteria. To address potential maternal inheritance, we cross-referenced bacterial OTUs from late stage (L6) and early stage (L1/L2) triatomines captured from the same nest. One nest from Arizona contained early and late stage *T. rubida*, as did one nest in Texas for *T. lecticularia.* In both instances there were common genera within all individuals (13 in *T. rubida*, 4 in *T. lecticularia*). We hypothesize that these shared taxa are the most likely candidates for vertical transmission.

While the significance of maternal inheritance is unclear, the effect of environmental bacteria is more evident. A prominent component of environmental microbe acquisition is potential vertebrate pathogens in the blood meal. In some hematophagous arthropods, vertebrate pathogens have evolved into symbionts (e.g. *Francisella* in the Gulf Coast tick [75]). Others, like sheep keds (*Melophagus ovinus* [83]) and a single kissing bug species (*Eratyrus mucronatus* [84]), were found to carry *Bartonella* species of an unknown phenotype. In our data, *Bartonella* was the second most abundant taxa found in every life stage of *T. rubida*. Molecular analysis showed that the bacterium is a pathogen acquired from *N. albigula*. A possible phenomenon for future consideration is whether *Bartonella* is a transcriptionally active component of *Triatoma* microbiomes, or a transient taxon reacquired with each feeding. *Mycobacterium*, the sixth most abundant OTU within *T. rubida*, provides another potential example of a pathogen [85, 86] acquired from the environment. Strict vertical inheritance and environmental uptake (horizontal transmission) are biologically distinct modes of acquiring bacterial symbionts. Triatomines engage in accessory feeding behaviours that potentially interconnect these two sources, such as coprophagy and kleptohematophagy [2, 45]. For instance, coprophagy is employed by first instar *Rhodnius prolixus* to acquire *Rhodococcus rhodnii* from parental faeces [87]. This form of symbiont acquisition is not strict maternal inheritance, because offspring do not acquire *Rhodococcus* in *utero* or from the mother’s ovaries. Instead, coprophagous symbiont acquisition represents both ‘indirect’ vertical transmission and environmental acquisition. Currently, we cannot determine whether *Triatoma* microbiome species-specificity is due to transmission of maternally provided bacteria or genetically determined uptake of environmental bacteria (e.g. the lack of *Dietzia* in *T. gerstaeckeri* and *T. sanguisuga* could be linked to the host’s close phylogenetic relationship), or a combination of both.

### 4.3 Dominant taxa and endosymbiosis

The consensus for many arthropod vectors is strong reliance on obligate bacterial endosymbionts (e.g. *Wigglesworthia* in *Glossina, Riesia* in *Pediculus* lice, *Coxiella-*like symbiont in ticks [16–20]) that facilitate essential functions like vitamin synthesis and participate in blood meal breakdown. Triatomines appear to establish less intimate symbioses with extracellular bacteria in their gut lumen, instead of possessing obligate intracellular symbionts [39, 42, 88]. Some studies indicated that *Rhodococcus* (an extracellular symbiont) was important for successful development and reproduction of *Rhodnius prolixus* [89–91]. However, later molecular studies showed *Rhodococcus* is not omnipresent throughout Triatominae, and not even among other *Rhodnius* species [43, 92]. In our results, *Dietzia*, a bacterium closely related to *Rhodococcus* [93], is the dominant bacterium in late instars of *T. rubida, T. protracta*, and *T. lecticularia. Dietzia* has been described from other triatomine species [33, 35, 36, 42–44] and other hematophagous insects (*Aedes albopictus* [94], *Glossina pallidipes* [95]), suggesting it may be an important mutualist. However, unlike typical primary symbionts, *Dietzia* does not seem to be transmitted vertically. In contrast to its’ obvious dominance in later instars, the presence of *Dietzia* in L1 is questionable. The ‘*decontam*’ dataset showed that five *T. rubida* L1s had 1-3 reads of *Dietzia* (from a median average of 1864 reads). Such low read numbers cannot be discriminated from marginal well-to-well contamination and do not provide evidence of *Dietzia* presence in first instars. Further studies with quantitative and *in situ* approaches are required to unequivocally determine the transmission mechanism and presence of *Dietzia* in first instar triatomines. One hypothesis is that individual bugs acquire *Dietzia* from other triatomines via ‘indirect’ vertical transmission (analogous to the *R. prolixus* and *Rhodococcus* example described above [87]) or from the environment, strategies that have been found in other true bug (Heteroptera) species, including trophallaxis, egg smearing, and endosymbiont reacquisition from soil [96–102]. To further investigate transmission and function of triatomine microbiomes, we will require tissue specific whole genome sequencing and functional transcriptomic studies.

### 4.4 Consistency of the patterns: biology vs. methodology

Previous studies on triatomine microbiomes have provided inconsistent and mutually conflicting results. They have suggested various factors, including ontogeny [38, 42], species identity [36, 40, 42, 43, 92], sex [34, 40], blood meal source [34], and *T. cruzi* infection [23, 35, 39, 40], as determinants of microbiome composition, whilst another study claimed triatomine microbiomes have no determining factor [33]. Since many were based on limited sample size (e.g. N=4 in [33]; N=14 in [34]; N=20 in [37]; N=9 in [36]; N=29 in [42]) and largely fragmented by host taxonomy, ontogeny, geographic origin, *T. cruzi* infection status, restricted to colony-reared bugs, and employed different molecular methods, it is difficult to draw comparative conclusions. We thus paid particular attention to our sampling design and molecular approach, ensuring that our study enabled multiple comparisons at different scales (i.e. different species from the same locality, different species from the same microhabitat, and different localities for the same species) across all ontogenetic stages. Furthermore, by introducing a novel method with a blocking primer (see methods section 2.4), we achieved greater precision for profiling microbiomes. As a result, this study presents multiple deterministic patterns consistent across several triatomine species for the first time.

Some of our findings contradict the patterns reported by other authors. The most conspicuous example is the ontogenetic decrease of microbiome diversity in North American species, which is supported by Mann et al. [40], but contrasts two studies on South American species [38, 43]. Oliveira et al. [38], reported an increase in microbiome diversity throughout ontogenetic development in *T. sordida* and Waltmann et al. [43] found no ontogenetic effect in *T. infestans.* There are two possible reasons for these differences. One is biological, because the other studies worked with South American species and species-specific differences are a clear component of microbiome dissimilarity, as our results and the results of others show [36, 42, 92]. The other is methodological, since the design of Oliveira et al.’s [38] study involved pooled samples and therefore does not allow evaluation of individual microbiome composition in different ontogenetic stages. Considering the among-individual compositional variability we observed, it is clear that pooling samples may have significantly distorted the profiles. A similar methodological artefact has been shown in mosquitoes [103]. The lack of ontogenetic differences in Waltmann et al. [43] could reflect the sample source (faeces) and incompleteness of the ontogenetic spectrum (L3 to adults only) rather than a real biological pattern in natural populations.

The importance of sampling the complete ontogenetic spectrum is well demonstrated by comparing our results with the recent survey of Mann et al. [40], which focused on *T. sanguisuga* and *T. gerstaeckeri*. By profiling microbiomes of 74 specimens, they also revealed a high degree of among-individual variability. However, since their study was based solely on adults, they reported weak species-specificity, whereas we found that species-specific microbiome patterns were more pronounced in early instars. In addition, Mann et al. [40] found support for locality-based effects which corroborates our findings for *T. rubida* from multiple locations in southern Arizona. These examples show that there is some consistency across triatomine microbiome studies, at least regarding US species. However, the current paucity of data does not allow for broader cross-study comparisons. To reach more generalisable conclusions for all Triatominae, we require added breadth (more studies) and depth (metagenomics and transcriptomics) of molecular data.

## 5. Conclusion

This study has contributed key information on triatomine microbiomes, which constitutes a crucial component of their biology. We identified ontogenetic shift, species identity, and the environment as the major factors determining microbiome composition in natural populations of *T. rubida, T. protracta, T. lecticularia, T. sanguisuga*, and *T. gerstaeckeri*, thus observing consistent deterministic patterns across multiple triatomine species for the first time. We hypothesise that the high among-individual variability of Triatominae microbiome assemblages is produced by inconsistent uptake of environmental bacteria, including vertebrate pathogens, and multiple indirect bacterial transmission strategies. The epidemiological relevance of Triatominae and their microbiome communities both warrant more in-depth exploration for successful implementation of microbiome-based vector control strategies. To achieve this, we advocate that future studies are designed to allow comparison of detected patterns across different triatomine populations, species, biogeographic areas, and environments.

## 6. Declarations

### Ethics approval

not applicable.

## Consent for publication

not applicable.

## Availability of data and materials

The datasets supporting the conclusions of this article and are included with the article and its additional files. A representative subsample of *cytB* sequences for all *Triatoma* species in this study were uploaded to NCBI (accession number: MT239320- MT239329). Representative sequences for *Bartonella vinsonii* are available under the following accession numbers: MT112947-MT112949. Raw, demultiplexed microbiome data were uploaded to ENA (project number: PRJEB36515). Sample metadata and R code is available as additional files to this manuscript.

## Competing interests

The authors declare that they have no competing interests.

## Funding

This research was funded by the Czech Science Foundation, grant no. 31-18-24707Y to E. Nováková. The funding body played no part in the study design, collection, analysis, an interpretation of data, or the writing of the manuscript.

## Author Contributions

EN designed the research. JJB, EN, SRR, WR collected the field samples. JOS provided supplementary samples. EN and JZ designed the novel 18S rRNA gene blocking primers. JZ, SRR, EN performed the DNA isolation and library preparation. AP performed *T. cruzi* screening. GB performed *Bartonella* sequencing. JJB, EN, SRR, AP, VH analysed the data and interpreted results. JJB, EN, SRR, VH wrote the manuscript. All authors contributed to improving the manuscript.

## Acknowledgements

We wish to thank Allison Hansen, Patrick Degnan, and Robert L. Smith for all their invaluable help and support in the USA. This research was performed with contributions from the Texas Park and Wildlife Management Department and the Bureau of Land Management (Gila District Office, Tucson, AZ).

## Additional Files

**Additional File 1:** Sample metadata including list of discarded contaminants.

**Additional File 2:** Results of ordination analyses based on ‘*basic’* and ‘*decontam’* datasets.

**Additional File 3:** Sample set phylogenetic background inferred from *coxB* sequences.

**Additional File 4:** Microbiome ontogenetic shift in other *Triatoma sp.*

**Additional File 5:** Significant difference in beta dispersion of the instar range groups (L1-L3 and L4-L6) calculated from the *ultraclean* dataset.

**Additional File 6:** NMDS analyses of *T. rubida* microbiomes from early instar (L1-L3) individuals found in different *N. albigula* nests at UADS.

**Additional File 7:** *Bartonella* phylogenetic analysis of gltA sequences retrieved from *T. rubida*.

**Additional File 8:** R code used in this study.

## Notes

### Competing Interest Statement

The authors have declared no competing interest.

