## Additional File 2 for "Ontogeny, species identity and environment dominate microbiome dynamics in wild populations of kissing bugs (Triatominae)"

**Additional File 2: Results of ordination analyses based on '*basic'* and '*decontam'* datasets.**

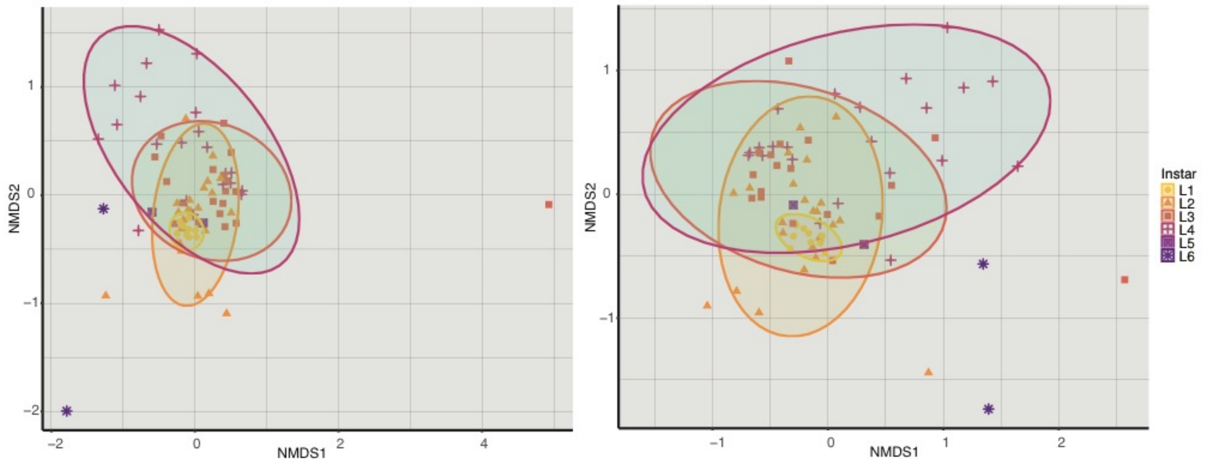


**A (Supplementing Figure 1C):** NMDS based on Bray-Curtis dissimilarity matrix calculated for *Triatoma rubida* ontogeny from the '*basic*' dataset (left) and '*decontam'* dataset (right). Each colour and shape correspond to a different instar; L6 stands for adults. Ellipses are statistically significant at the confidence interval of 0.05.

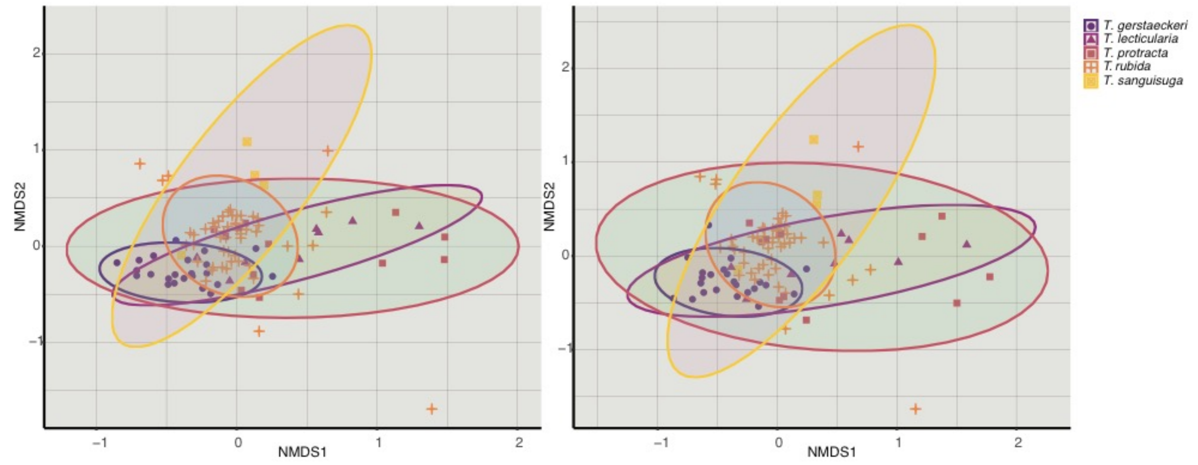


**B (Supplementing Figure 3):** Microbiome species-specific differences. NMDS ordination for the early instar dataset (L1-L3) of five *Triatoma* species based on Bray-Curtis dissimilarities calculated from from the '*basic*' dataset (left) and '*decontam'* dataset (right).

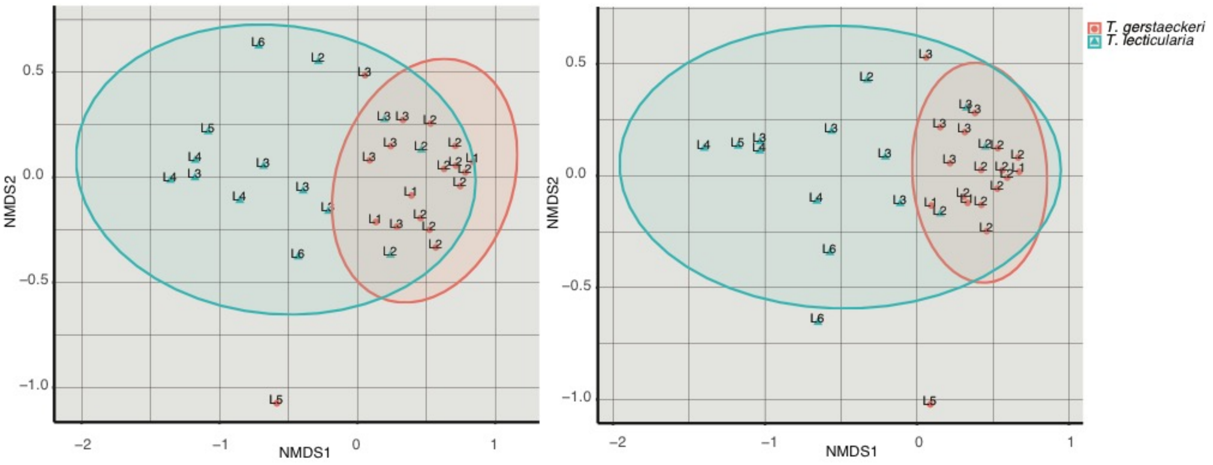


**C (Supplementing Figure 4):** Non-metric multidimensional scaling analysis for for *Triatoma gerstaeckeri* and *Triatoma lecticularia* from the same nest in Chaparral, TX; calculated from from the '*basic*' dataset (left) and '*decontam'* dataset (right).

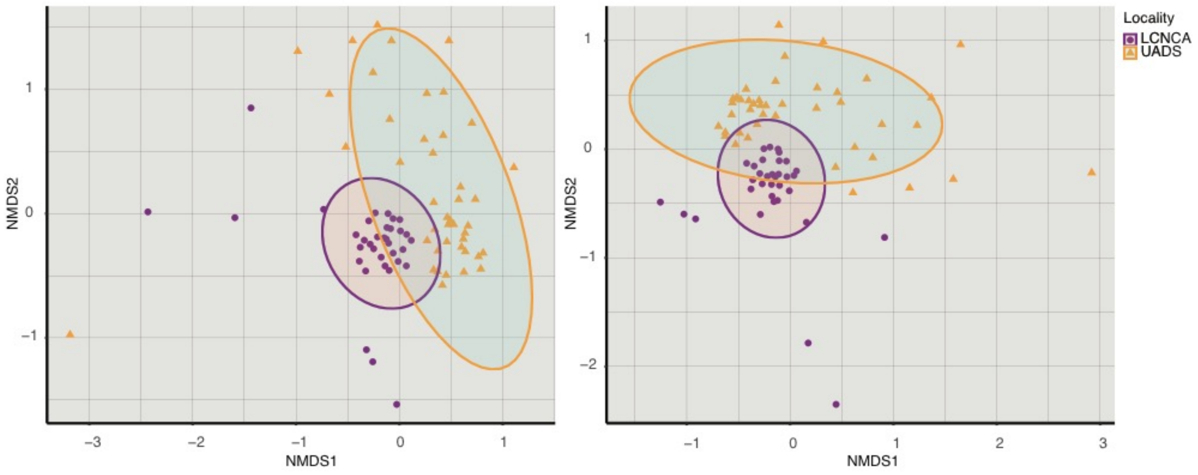


**D (Supplementing Figure 6):** Non-metric multidimensional scaling analysis for *T. rubida* collected from two locations in southern Arizona. LCNCA stands for Las Cienegas National Conservation Area, UADS for University of Arizona Desert Station; calculated from from the '*basic*' dataset (left) and '*decontam'* dataset (right).
