## Additional File 3 for "Ontogeny, species identity and environment dominate microbiome dynamics in wild populations of kissing bugs (Triatominae)"

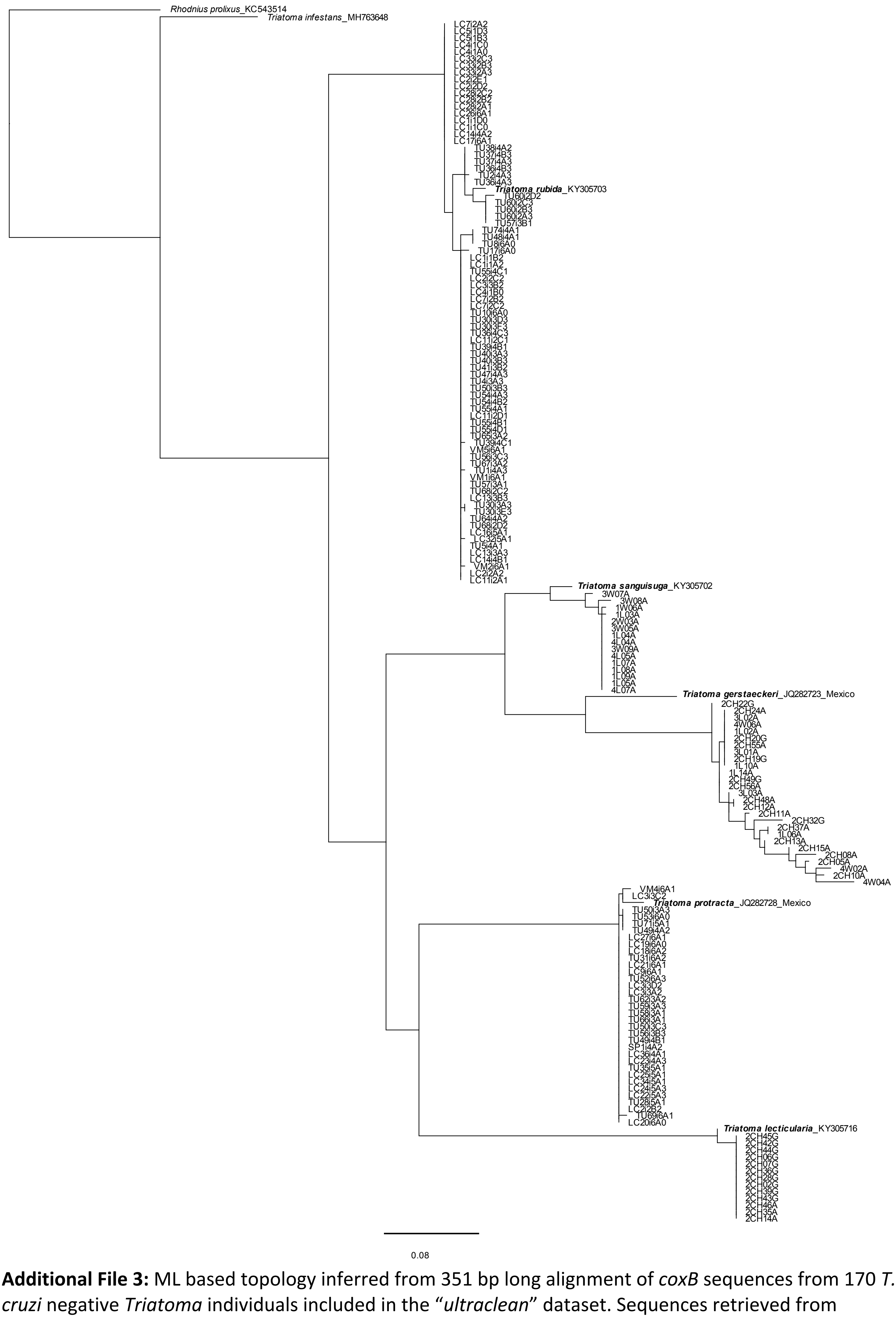


**Additional File 3:** ML based topology inferred from 351 bp long alignment of *coxB* sequences from 170 *T. cruzi* negative *Triatoma* individuals included in the "*ultraclean*" dataset. Sequences retrieved from GenBank are designated with the accession numbers.

(Rhodnius_prolixus_KC543514:0.12680117000000002,('Triatoma
_infestans_MH763648':0.08185486999999991,((LC7i2A2:0.0,((LC5i1D3:0.0,(LC5i1B3:0.0,(LC4i1C0:0.0,(LC4i
1A0:0.0,(LC33i2C3:0.0,(LC33i2B3:0.0,(LC33i2A3:0.0,(LC2i2E1:0.0,(LC2i2D2:0.0,(LC28i2C2:0.0,(LC28i2B2:0.0,(
LC28i2A1:0.0,(LC26i6A1:0.0,(LC1i1D0:0.0,(LC1i1C0:0.0,(LC14i4A2:0.0,LC17i6A1:0.0):0.0):0.0):0.0):0.0):0.0):
0.0):0.0):0.0):0.0):0.0):0.0):0.0):0.0):0.0):0.0):0.0,(((TU38i4A2:0.0,(TU37i4B3:0.0,(TU37i4A3:0.0,(TU36i4B3:
0.0,(TU2i4A3:0.003341380000000005,TU36i4A3:0.0):0.0):0.0):0.0):0.0):0.0,(Triatoma_rubida_KY305703:0.
010401959999999988,(TU60i2D2:0.006910110000000025,(TU60i2C3:0.0,(TU60i2B3:0.0,(TU60i2A3:0.0,TU
57i3B1:0.0):0.0):0.0):0.0):0.010668310000000014):0.006887609999999885):0.010197080000000025,((T
U74i4A1:0.0,(TU48i4A1:0.0,TU8i6A0:0.0):0.0):0.010151819999999978,(TU17i6A0:0.006715230000000016
5,(LC1i1B2:0.0,(LC1i1A2:0.0,(TU55i4C1:0.0,((LC2i2C2:0.0,(LC3i3B2:0.0,(LC4i1B0:0.0,(LC7i2B2:0.0,(LC7i2C2:0
.0,(TU10i6A0:0.0,(TU30i3D3:0.0,(TU30i3F3:0.0,(TU36i4C3:0.0,(LC11i2C1:0.0,(TU39i4B1:0.0,(TU40i3A3:0.0,(
TU40i3B3:0.0,(TU41i3B2:0.0,(TU47i4A3:0.0,(TU4i3A3:0.0,(TU50i3B3:0.0,(TU54i4A3:0.0,(TU54i4B2:0.0,(TU5
5i4A1:0.0,(LC11i2D1:0.0,(TU55i4B1:0.0,((TU55i4D1:0.0,(TU65i3A2:0.0,(TU39i4C1:0.003352140000000035
,VM5i6A1:0.0):0.0):0.0):0.0,((TU56i3C3:0.0,(TU67i3A2:0.0,(TU1i4A3:0.003335440000000023,VM1i6A1:0.0)
:0.0):0.0):0.0,((TU57i3A1:0.0,(TU68i2C2:0.0,(LC13i3B3:0.0,(TU30i3A3:0.0,TU30i3E3:0.0):0.00333583999999
9979):0.0):0.0):0.0,((TU64i4A2:0.0,(TU68i2D2:0.0,(LC16i5A1:0.0,LC32i5A1:0.003357090000000007):0.0):0.
0):0.0,(TU5i4A1:0.0,(LC13i3A3:0.0,(LC14i4B1:0.0,VM2i6A1:0.003361159999999974):0.0):0.0):0.0):0.0):0.0):
0.0):0.0):0.0):0.0):0.0):0.0):0.0):0.0):0.0):0.0):0.0):0.0):0.0):0.0):0.0):0.0):0.0):0.0):0.0):0.0):0.0):0.0):0.0
,(LC2i2A2:0.0,LC11i2A1:0.0):0.0):0.0):0.0):0.0):0.0):0.0):0.006736799999999987):0.006878080000000009):
0.0):0.09744577999999998,((('Triatoma|_sanguisuga_KY305702':0.01824619999999999,(3W07A:0.00601
50999999999955,(3W08A:0.010262140000000003,(1W06A:0.003230080000000024,(1L03A:0.0032121600
000000194,(2W03A:0.0,(3W05A:0.0,(1L04A:0.0,(4L04A:0.0,(3W09A:0.0,(4L05A:0.0,(1L07A:0.0,(1L08A:0.0,(
1L09A:0.0,(1L05A:0.0,4L07A:0.0):0.0):0.0):0.0):0.0):0.0):0.0):0.0):0.0):0.0):0.0):0.00288346999999999
9):0.011071439999999988):0.029484420000000001):0.03807145000000001,(Triatoma_gerstaeckeri_JQ282
723_Mexico:0.07655411000000001,(2CH22G:0.0,((((2CH24A:0.0,3L02A:0.0):0.0,(4W06A:0.0,(1L02A:0.0,(2
CH20G:0.0,(2CH55A:0.0,3L01A:0.0):0.0):0.0):0.0):0.0):0.0,(2CH19G:0.0,1L10A:0.0):0.0):0.00438141000000
003,(1L14A:0.0,((2CH49G:0.0,2CH56A:0.0):0.0,(3L03A:0.0,((2CH48A:0.0,2CH12A:0.0):0.00400949999999999
99,(2CH11A:0.003304019999999963,(2CH32G:0.023743500000000003,((2CH37A:0.0,1L06A:0.0):0.0068855
50000000018,(2CH13A:0.0,(2CH15A:0.0,(2CH08A:0.016816449999999983,(2CH05A:0.00287369999999999
79,(4W02A:0.012423799999999985,(2CH10A:0.006612630000000008,4W04A:0.03180011000000005):0.0
):0.008938769999999985):0.008063719999999996):0.005761789999999989):0.020420760000000038):0.
004101429999999615):0.004776070000000021):0.007496929999999957):0.013673909999999956):0.0):
0.008049729999999977):0.0):0.0):0.006282140000000047):0.10615423999999996):0.06776886999999999
8):0.09992028999999997,((((VM4i6A1:0.006552000000000002,(LC3i3C2:0.0,Triatoma_protracta_JQ28272
8_Mexico:0.017509760000000004):0.0):0.003252030000000724,(TU50i3A3:0.0,(TU53i6A0:0.0,(TU71i5A1:
0.0,TU49i4A2:0.0):0.0):0.0):0.003253530000000006):0.0,(LC27i6A1:0.0,(LC19i6A0:0.0,(LC18i6A2:0.0,(TU31i6
A2:0.0,(LC21i6A1:0.0,(LC9i6A1:0.0,(TU52i6A3:0.0,(LC3i3D2:0.0,(LC3i3A2:0.0,(TU62i3A2:0.0,(TU59i3A3:0.0,(
TU58i3A1:0.0,(TU66i3A1:0.0,(TU50i3C3:0.0,(TU56i3B3:0.0,(TU49i4B1:0.0,(SP1i4A2:0.0,(LC36i4A1:0.0,(LC23
i4A3:0.0,(TU35i5A1:0.0,(LC25i5A1:0.0,(LC34i5A1:0.0,(LC24i5A3:0.0,(LC22i5A3:0.0,(TU28i5A1:0.0,(LC2i2B2:0
.0,(TU69i6A1:0.006536070000000005,LC20i6A0:0.0):0.0):0.0):0.0):0.0):0.0):0.0):0.0):0.0):0.0):0.0):0.0):0.0)
:0.0):0.0):0.0):0.0):0.0):0.0):0.0):0.0):0.0):0.0):0.0):0.0):0.0):0.0):0.0):0.16803601,(Triatoma_lecticularia_KY
305716:0.0,(2CH45G:0.0,(2CH42G:0.0,(2CH44G:0.0,(2CH06G:0.0,(2CH07G:0.0,(2CH36G:0.0,(2CH28G:0.0,(2
CH02G:0.0,(2CH39G:0.0,(2CH43G:0.0,(2CH46A:0.0,(2CH35A:0.0,2CH14A:0.0):0.0):0.0):0.0):0.0):0.0):0.0):0.
0):0.0):0.0):0.0):0.0):0.015897889999999997):0.25078821000000007):0.027690150000000024):0.0482790
90000000024):0.14163196999999994):0.12680117000000002);
