## Additional File 4 for "Ontogeny, species identity and environment dominate microbiome dynamics in wild populations of kissing bugs (Triatominae)"

**Additional File 4: Microbiome ontogenetic shift in other *Triatoma sp.***

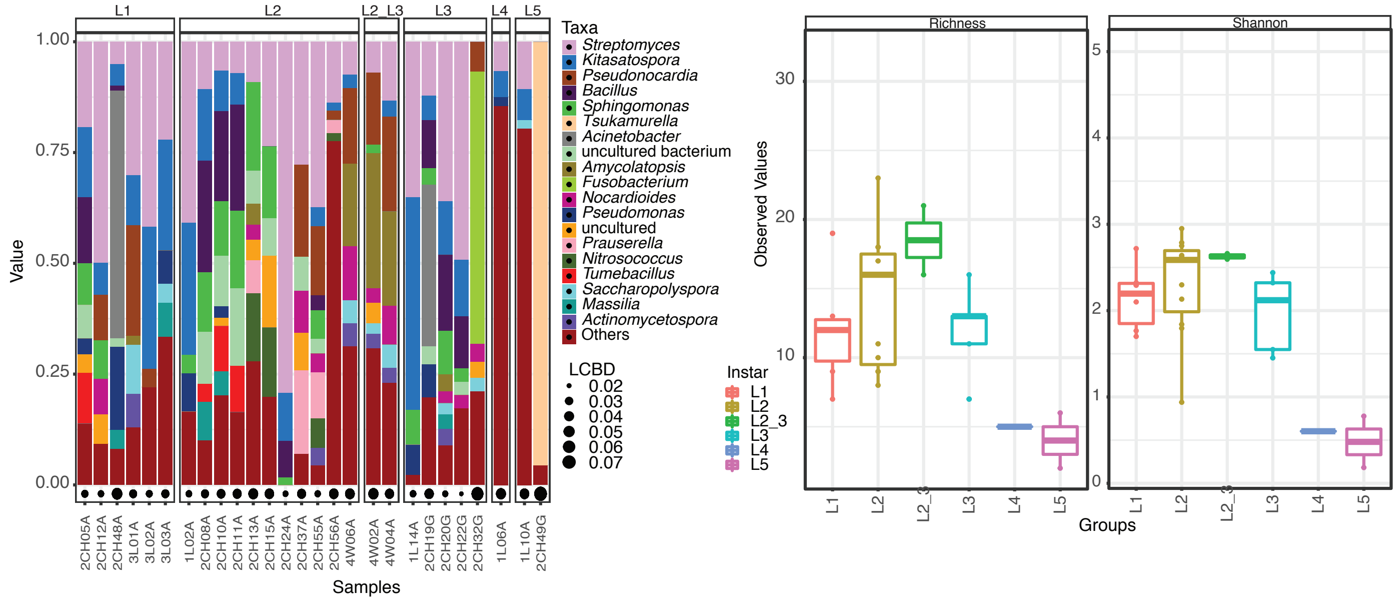


**A (Supplementing Figure 1B)**: Microbiome profiles (top 20 bacterial genera) and diversity indices for the ontogenetic range of *Triatoma gerstaeckeri* based on the '*ultraclean*' dataset.

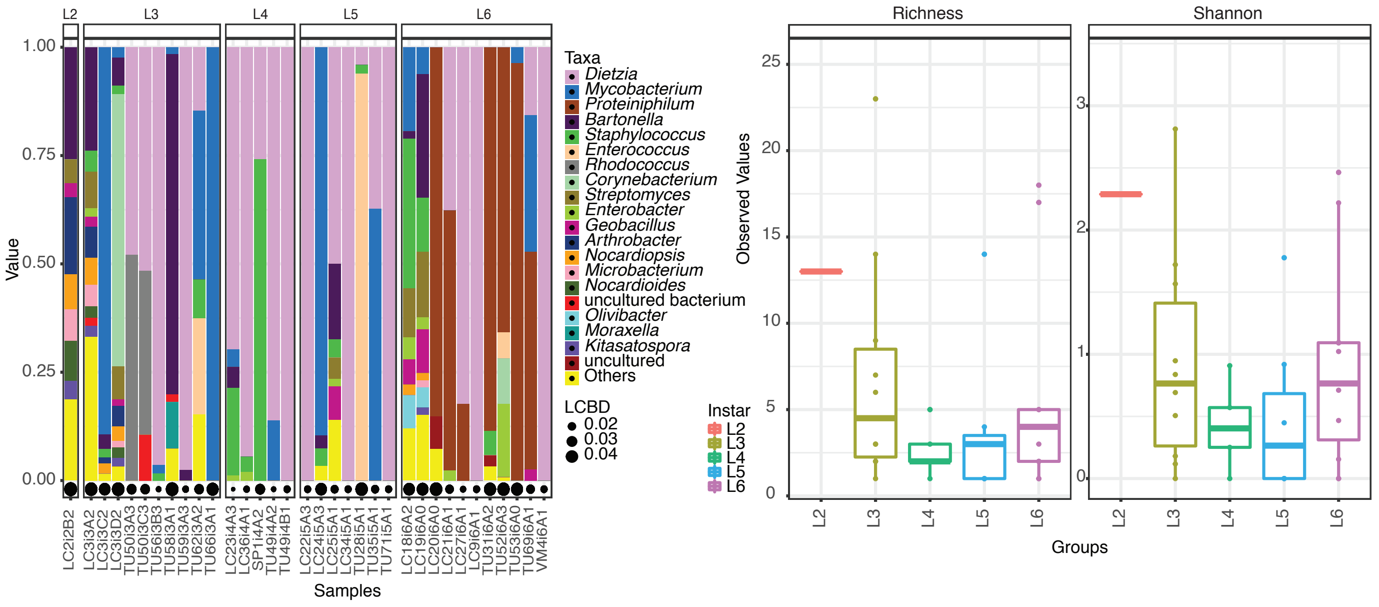


**B (Supplementing Figure 1B)**: Microbiome profiles (top 20 bacterial genera) and diversity indices for the available ontogenetic range of *Triatoma protracta* based on the '*ultraclean*' dataset. Pink denotes *Dietzia* which displays a similar cumulative pattern to *T. rubida* discussed in the main manuscript.

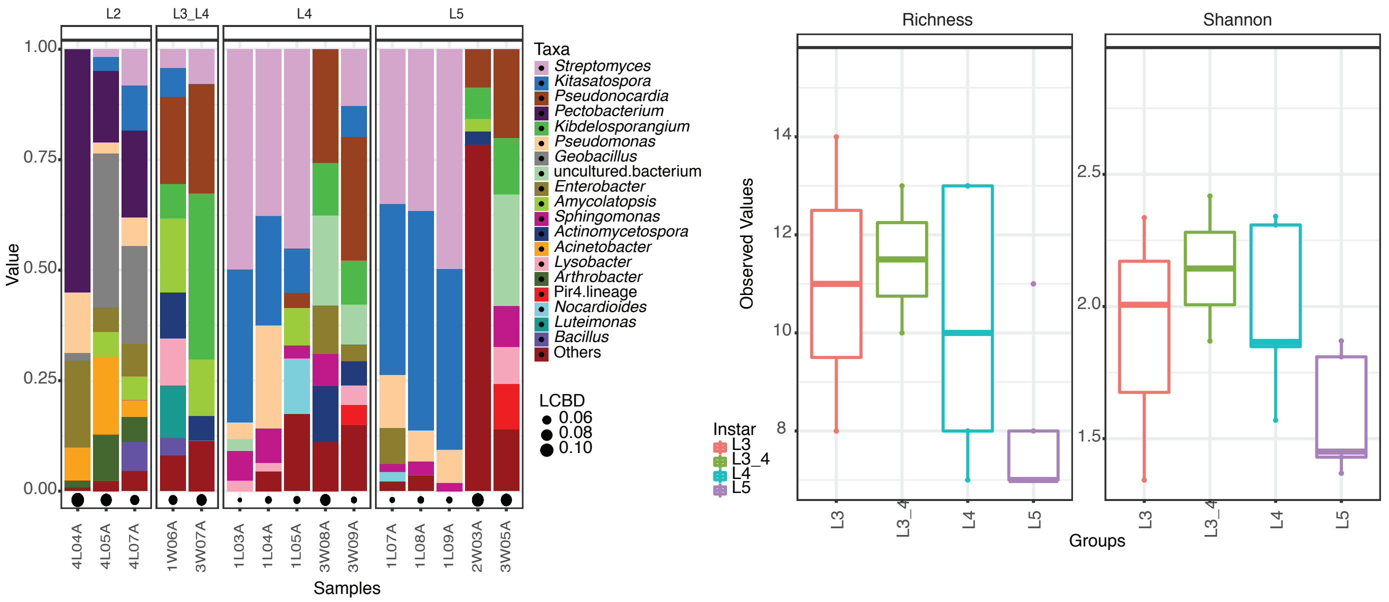


**C (Supplementing Figure 1B)**: Microbiome profiles (top 20 bacterial genera) and diversity indices for the available ontogenetic range of *Triatoma sanguisuga* based on the '*ultraclean*' dataset.

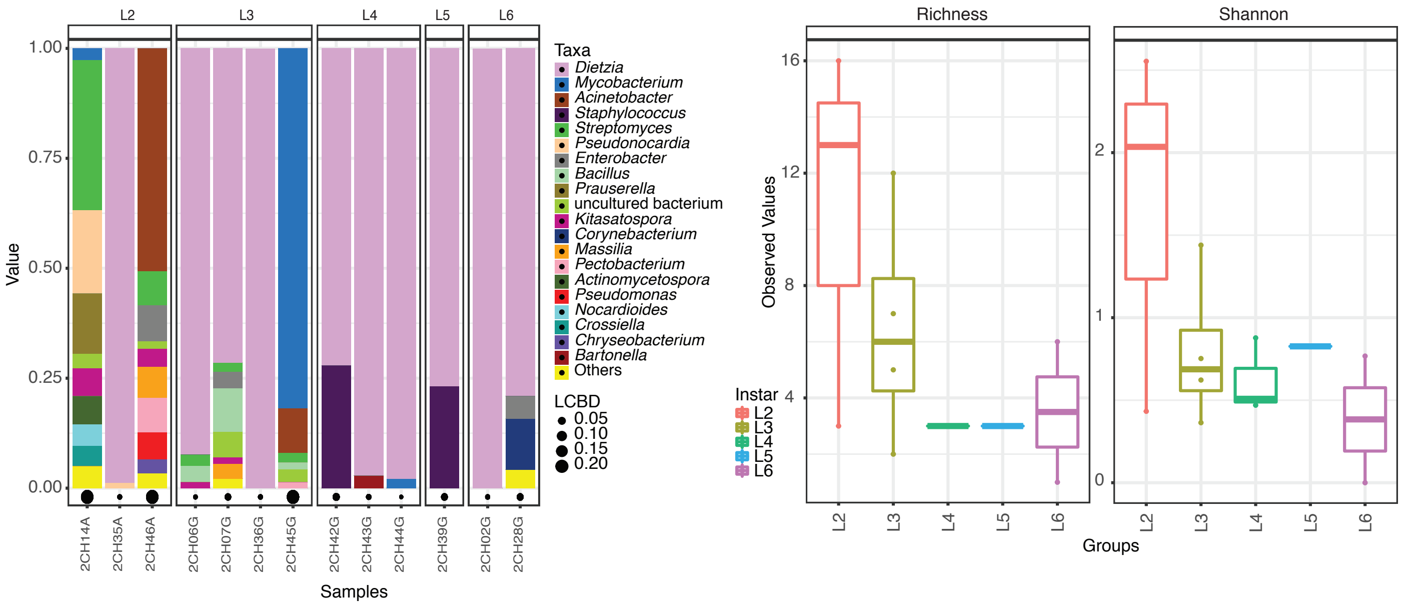


**D (Supplementing Figure 1B)**: Microbiome profiles (top 20 bacterial genera) and diversity indices for the available ontogenetic range of *Triatoma lecticularia* based on the '*ultraclean*' dataset. *T. lecticularia* is the third (after *T. rubida* and *T. protracta*) of our study species to display a similar pattern of *Dietzia* over ontogenetic development.
