## Additional File 5 for "Ontogeny, species identity and environment dominate microbiome dynamics in wild populations of kissing bugs (Triatominae)"

| | Df | Sum | Sq Mean | Sq F value | Pr(>F) |
|---|---|---|---|---|---|
| Groups | 1 | 0.5625 | 0.56249 | 11.749 | 0.0007677 *** |
| Residuals | 166 | 7.9475 | 0.04788 | | |

Signif. codes:  0 '***' 0.001 '**' 0.01 '*' 0.05 '.' 0.1 ' ' 1

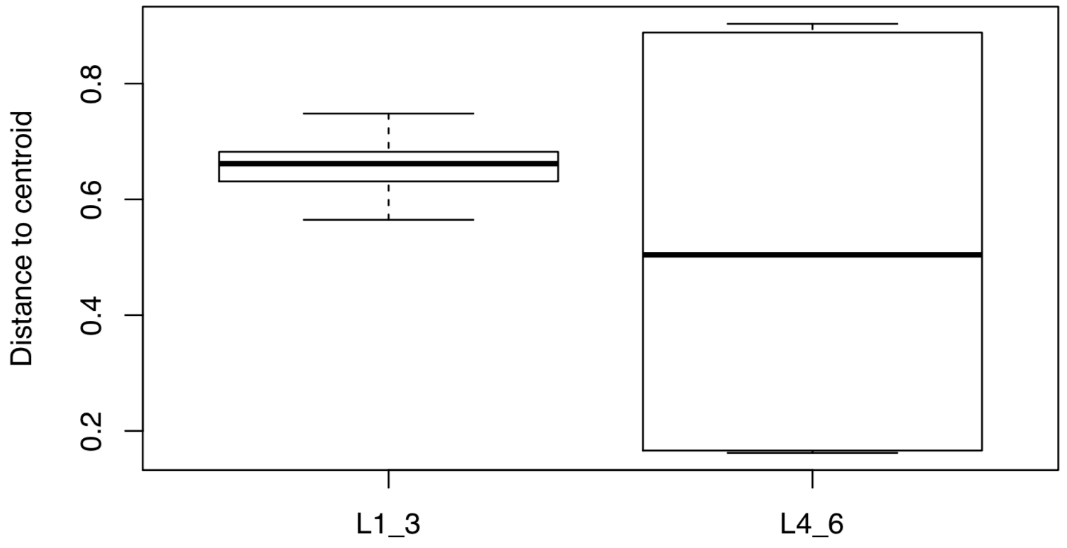


**Additional File 5:** Significant difference in beta dispersion of the instar range groups (L1-L3 and L4-L6) calculated from the *ultraclean* dataset. The beta dispersion analyses were performed with Jaccard index distance matrix using Vegan package (v2.5.6) in R software (v3.6.1).
