## Additional File 6 for "Ontogeny, species identity and environment dominate microbiome dynamics in wild populations of kissing bugs (Triatominae)"

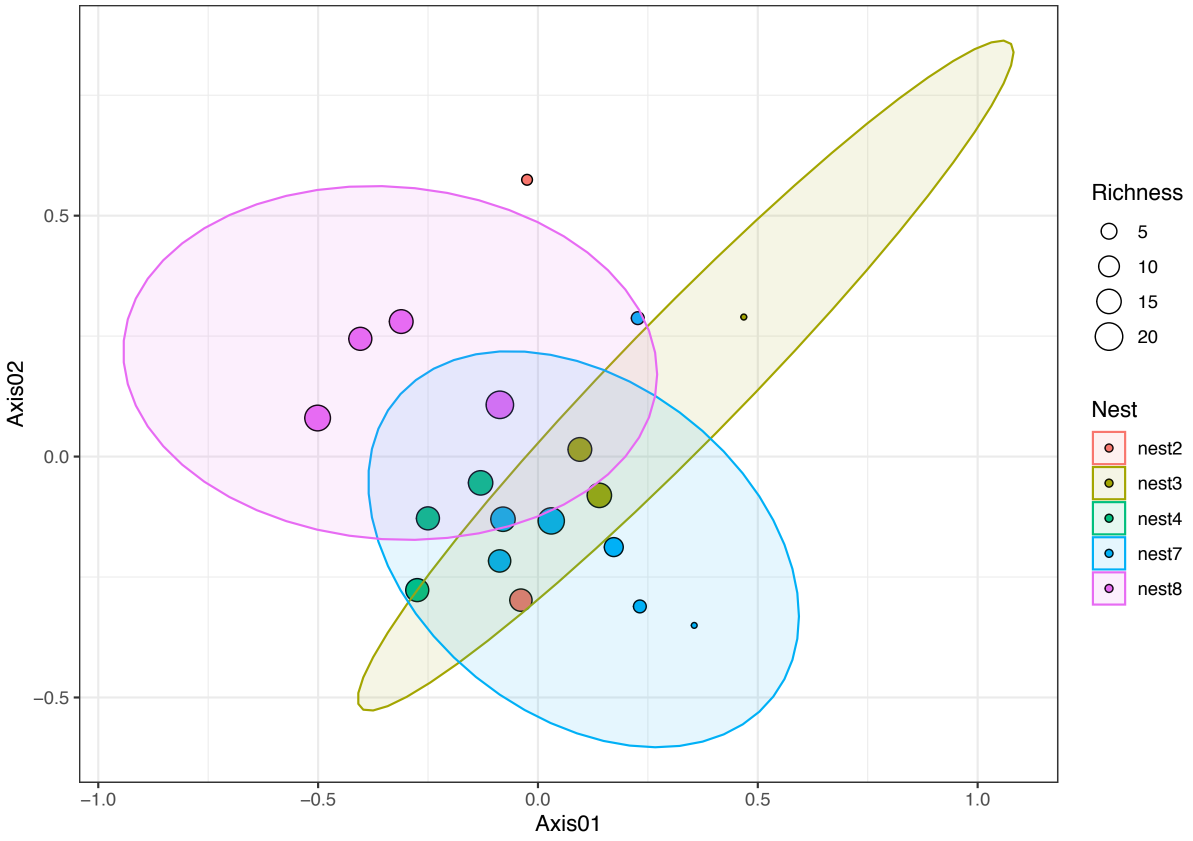


**Additional File 6:** NMDS analyses of *T. rubida* microbiomes from early instar (L1-L3) individuals found in different *N. albigula* nests found at UADS.
