## Additional File 7 for "Ontogeny, species identity and environment dominate microbiome dynamics in wild populations of kissing bugs (Triatominae)"

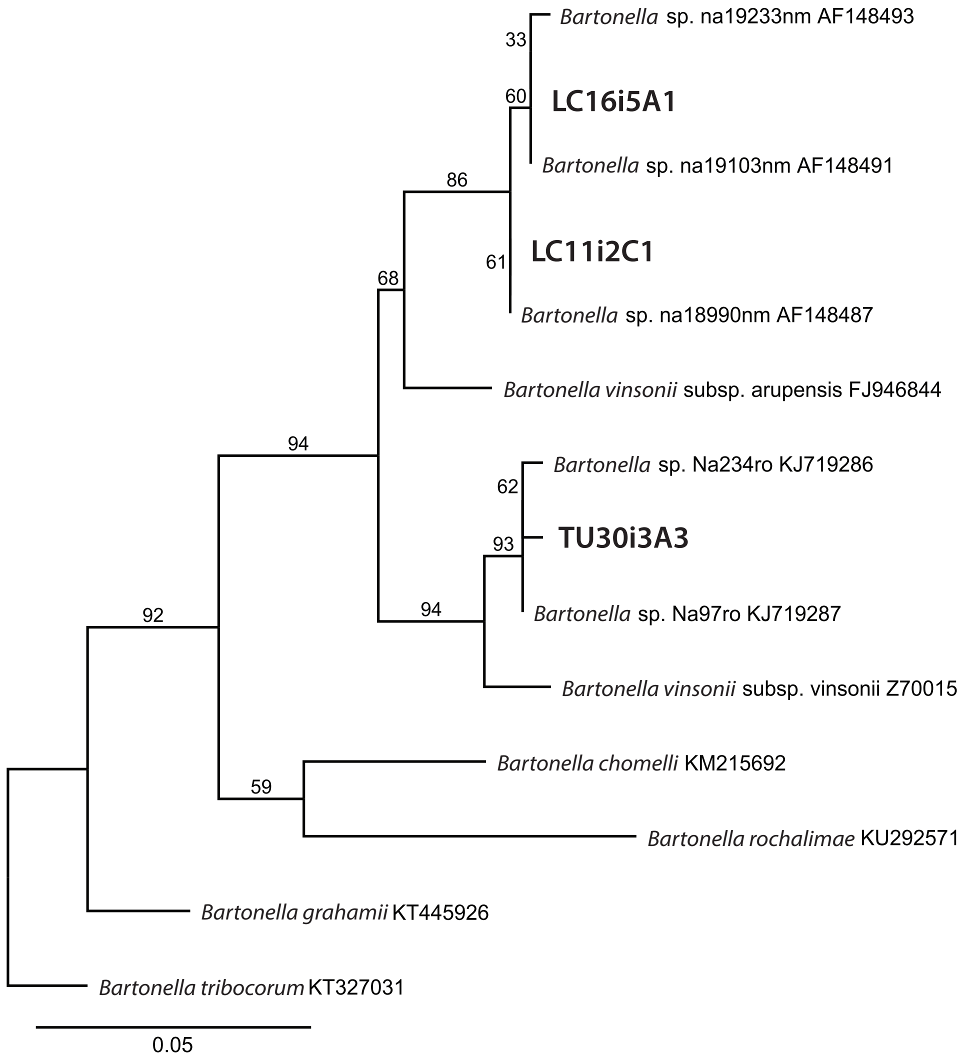


**Additional File 7.** Maximum-likelihood phylogenetic tree for the partial *gltA* sequence of *Bartonella* spp. Designated samples represent 12 (LC16i5A1), 5 (LC11i2C1) and 1 (TU30i3A3) other sequences retrieved from *T. rubida* individuals in this study. The numbers at the nodes designate bootstrap values.
