## Additional File 8 for "Ontogeny, species identity and environment dominate microbiome dynamics in wild populations of kissing bugs (Triatominae)"

---
title: "Additional_file_manuscript"
author: "Anbu Poosakkannu"
date: "6/1/2020"
output:
  pdf_document: default
  html_document: default
---

#script for removing potential contaminant from basic data #output is called as decontam data set

```{r, message=FALSE}

#load libraries
library(phyloseq); packageVersion("phyloseq")
library(ggplot2); packageVersion("ggplot2")
library(decontam); packageVersion("decontam")
library(dplyr); packageVersion("dplyr") #require to filter and clean up the
data
library(microbiome); packageVersion("microbiome") #require o write the output
files


#upload the files
#upload the OTU abundance file
abund_table<-
read.csv("otu_norarefaction_abund.csv",row.names=1,check.names=FALSE)

#transpose the OTU abundance data to have sample names on rows
abund_table<-t(abund_table)

#upload the meta data file
meta_table<-
read.csv("MapRunJune2019_updated30102019_complete.csv",row.names=1,check.names=FALSE)


#load the taxonomy
OTU_taxonomy<-
read.csv("otu_norarefaction_taxa.csv",row.names=1,check.names=FALSE)



#Convert the data to phyloseq format
OTU = otu_table(as.matrix(abund_table), taxa_are_rows = FALSE)
TAX = tax_table(as.matrix(OTU_taxonomy))
SAM = sample_data(meta_table)
physeq<-merge_phyloseq(phyloseq(OTU, TAX),SAM)
physeq
```

This document was created by an application that isn't licensed to use novaPDF.
Purchase a license to generate PDF files without this notice.

```
#column names of taxonomy file
colnames(tax_table(physeq))
#if need to be renamed #here i renamed Domain into Kingdom
colnames(tax_table(physeq)) <- c("Kingdom", "Phylum", "Class", "Order",
"Family", "Genus", "Species")

summarize_phyloseq(physeq)

#clean up and filtering the data

#removing the contaminants to keep only bacteria
physeq_clean <- physeq %>%
  subset_taxa(
    Kingdom == "Bacteria" &
    Kingdom  != "Archaea" &
    Kingdom  != "unclassified" &
    Family  != "mitochondria" &
    Class   != "Chloroplast"
  )

physeq_clean

#identify Contaminants by Frequency
contamdf.freq <- isContaminant(physeq_clean, method="frequency",
conc="postPCR_concentration")
head(contamdf.freq)

table(contamdf.freq$contaminant)

head(which(contamdf.freq$contaminant))

#plot contaminant taxa frequency Vs pcr product concentration
set.seed(100)
plot_frequency(physeq_clean, taxa_names(physeq_clean)
[sample(which(contamdf.freq$contaminant),117)], conc="postPCR_concentration")
+
    xlab("PCR product concentration (ng/µL)")


#filter the contaminant identified by decontaminant package
physeq_clean_decontam1 <- prune_taxa(!contamdf.freq$contaminant, physeq_clean)
physeq_clean_decontam1


#remove an additional OTU (unknown bacteria) present in negative samples
badtaxa = "OTU_1849"
allTaxa = taxa_names(physeq_clean_decontam1)
allTaxa <- allTaxa[!(allTaxa %in% badtaxa)]
physeq_clean_decontam2 = prune_taxa(allTaxa, physeq_clean_decontam1)

physeq_clean_decontam2
```

This document was created by an application that isn't licensed to use novaPDF.
Purchase a license to generate PDF files without this notice.

````
write_phyloseq(physeq_clean_decontam2, type = "OTU", path = getwd())

write_phyloseq(physeq_clean_decontam2, type = "TAXONOMY", path = getwd())


```




#script for the ultra clean data #prdouce Alpha diversity, coordination
analyses and taxonomy profile figures in main text


```{r, message=FALSE}

##################### 1.0 INTRODUCTION ######################
# Get BiocManager working, and relevant packages installed
if (!requireNamespace("BiocManager", quietly = TRUE))
  install.packages("BiocManager")
BiocManager::install(c("AnnotationDbi", "DESeq2", "GO.db", "impute",
"phyloseq",
                       "preprocessCore", "biomformat", "decontam",
"microbiome"))

library(biomformat) #needed for reading biom
library(phyloseq) #needed for proper microbiome analysis
library(vegan) #community ecology analyses
library(ggplot2) #graphs
library(ape) #require to load the tree
library(dplyr) #to filter and clean up the data
library(cowplot) #multiple ggplots
library(GUniFrac) #for unifrac distance based ordination
library(phangorn) #for unifrac distance based ordination using phylogenetic
tree
library(viridis) #colour package

#Read the OTU file
abund_table<-
read.csv("otu_ultraclean_abund.csv",row.names=1,check.names=FALSE)
#transpose the OTU abundance data to have sample names on rows
abund_table<-t(abund_table)
#Read in the complete meta data file
meta_table<-read.csv("mapNovember.csv",row.names=1,check.names=FALSE)
#Read the taxonomy
OTU_taxonomy<-
read.csv("otu_ultraclean_taxa.csv",row.names=1,check.names=FALSE)


colnames(meta_table)<-c("ReadCount", "Plate", "WellPosition", "Organism",
"State",
                        "Tcruzi","Host_Taxa", "Instar", "Instar_range", "Sex",
"Locality",
                        "Nest", "Latitude", "Longitude", "Sample")
````

This document was created by an application that isn’t licensed to use novaPDF.
Purchase a license to generate PDF files without this notice.

```
###  1.1 - Phyloseq adaptation + cleanup
#Convert the data to phyloseq format
OTU = otu_table(as.matrix(abund_table), taxa_are_rows = FALSE)
TAX = tax_table(as.matrix(OTU_taxonomy))
SAM = sample_data(meta_table)
physeq<-merge_phyloseq(phyloseq(OTU, TAX),SAM)

clean_data<-physeq
#column names of taxonomy file
colnames(tax_table(clean_data))

# Create some new subsets for later use here:
rubida_rarefy <- clean_data %>% subset_samples(Host_Taxa == "Trubida")
chaparral <- clean_data %>% subset_samples(Locality == "Chaparral")
tucson <- clean_data %>% subset_samples(Locality == "UADS" | Locality ==
"LCNCA")
young <- clean_data %>% subset_samples(Instar == "L1"|Instar == "L2"|Instar ==
"L3")


############## 1.2 UNCONSTRAINED ORDINATION ##############
# Set.seed for randomisation, making results reproducible
set.seed(5)
# 1.2.1 NMDS on rubida ontogeny
rubida_nmds <- ordinate(
  physeq = rubida_rarefy,
  method = "NMDS",
  distance = "bray"
)
## Stress = 0.15
## Plot it
rub2<-plot_ordination(
  physeq = rubida_rarefy,
  ordination = rubida_nmds,
  color = "Instar",
  shape = "Instar",
  title = "NMDS of Rubida ontogeny") +
  scale_color_viridis(option="plasma", discrete = TRUE, begin = 0.2, end =
0.9, direction = -1) +
  theme(panel.grid.major = element_line(size = 0.05, linetype = 'solid',
colour = "white"),
        panel.grid.minor = element_line(size = 0.05, linetype = 'solid',
colour = "white"),
        plot.background=element_rect(fill = "grey90"),
        panel.background = element_rect(fill = 'black'),
        axis.line = element_line(colour = "white"),) +
  geom_point(alpha = 1, size = 4) +
  stat_ellipse(geom='polygon', aes(colour=Instar, fill=Instar), alpha = 0.1)
print(rub2)
## 1.2.2 PERMANOVA ON ONTOGENY
rubida_bray <- phyloseq::distance(rubida_rarefy, method = "bray")
rubida_rarefy_df <- data.frame(sample_data(rubida_rarefy))
```

This document was created by an application that isn’t licensed to use novaPDF.
Purchase a license to generate PDF files without this notice.

```
adonis(rubida_bray ~ Instar, data = rubida_rarefy_df, permutations = 999,
strata = rubida_rarefy_df$Locality)
beta1 <- betadisper(rubida_bray, rubida_rarefy_df$Instar, type = "median",
bias.adjust = TRUE)
permutest(beta1, pairwise = TRUE, permutations = 999, model = "direct")



####### 1.2.3 NMDS on multi-species nest from Chaparral, TX
## Now NMDS
chap_nmds <- ordinate(
  physeq = chaparral,
  method = "NMDS",
  distance = "bray"
)
## 0.09 stress
chap2<-plot_ordination(
  physeq = chaparral,
  ordination = chap_nmds,
  color = "Host_Taxa",
  shape = "Host_Taxa",
  title = "NMDS of Chaparral Multi-Species Nest") +
  scale_color_viridis(option="plasma", discrete = TRUE, begin = 0.5, end =
0.8, direction = 1) +
  theme(panel.grid.major = element_line(size = 0.02, linetype = 'solid',
colour = "black"),
        panel.grid.minor = element_line(size = 0.02, linetype = 'solid',
colour = "black"),
        plot.background=element_rect(fill = "white"),
        panel.background = element_rect(fill = 'white'),
        axis.line = element_line(colour = "black"),) +
  geom_point(alpha = 1, size = 4) +
  geom_text(aes(label=Instar),hjust=1, vjust=0, size=3, colour = "black")+
  stat_ellipse(geom='polygon', aes(colour=Host_Taxa, fill=Host_Taxa), alpha =
0.1, level = 0.95)
print(chap2)
## 1.2.4 PERMANOVA ON CHAPARRAL
chap_bray <- phyloseq::distance(chaparral, method = "bray")
chap_df <- data.frame(sample_data(chaparral))
adonis(chap_bray ~ Host_Taxa, data = chap_df, permutations = 999, strata =
chap_df$Instar) #27% of the variation is species level
## Beta dispersion
beta2 <- betadisper(chap_bray, chap_df$Host_Taxa)
permutest(beta2, pairwise = TRUE)



## 1.2.5 NMDS on rubida from 2 locations in Arizona
tuc_nmds <- ordinate(
  physeq = tucson,
  method = "NMDS",
  distance = "bray"
)
## 0.17 stress
```

This document was created by an application that isn't licensed to use novaPDF.
Purchase a license to generate PDF files without this notice.

```
tuc1<-plot_ordination(
  physeq = tucson,
  ordination = tuc_nmds,
  color = "Locality",
  shape = "Instar_range",
  title = "NMDS of Tucson rubida") +
  scale_color_viridis(option = "plasma", discrete = TRUE, begin = 0.3, end =
0.8)+
  theme(panel.grid.major = element_line(size = 0.02, linetype = 'solid',
colour = "black"),
        panel.grid.minor = element_line(size = 0.02, linetype = 'solid',
colour = "black"),
        plot.background=element_rect(fill = "gray90"),
        panel.background = element_rect(fill = 'white'),
        axis.line = element_line(colour = "black"),) +
  geom_point(alpha = 1, size = 4) +
  #geom_text(aes(label=Instar),hjust=1, vjust=0, size=3, colour = "black")+
  stat_ellipse(geom='polygon', aes(colour=Locality, fill=Locality), alpha =
0.1, level = 0.95)
print(tuc1)
## 1.2.6. PERMANOVA ON TUCSON NEST RUBIDA
tuc_bray <- phyloseq::distance(tucson, method = "bray")
tuc_df <- data.frame(sample_data(tucson))
adonis(tuc_bray ~ Locality*Instar, data = tuc_df)
beta3 <- betadisper(tuc_bray, tuc_df$Locality)
permutest(beta3)



######  1.2.7 NMDS on young instar species-specific differences
young_nmds <- ordinate(
  physeq = young,
  method = "NMDS",
  distance = "bray"
)
### stress = ~0.2
young1<-plot_ordination(
  physeq = young,
  ordination = young_nmds,
  color = "Host_Taxa",
  shape = "Host_Taxa",
  title = "Young instars") +
  scale_color_viridis(option = "plasma", discrete = TRUE, begin = 0.2, end =
0.9)+
  theme(panel.grid.major = element_line(size = 0.02, linetype = 'solid',
colour = "black"),
        panel.grid.minor = element_line(size = 0.02, linetype = 'solid',
colour = "black"),
        plot.background=element_rect(fill = "white"),
        panel.background = element_rect(fill = 'white'),
        axis.line = element_line(colour = "black"),) +
  geom_point(alpha = 1, size = 4)+
  stat_ellipse(geom='polygon', aes(colour=Host_Taxa, fill=Host_Taxa), alpha =
0.1, level = 0.95)
```

This document was created by an application that isn’t licensed to use novaPDF.
Purchase a license to generate PDF files without this notice.

```
print(young1)
## 1.2.8
young_bray <- phyloseq::distance(young, method = "bray")
young_df <- data.frame(sample_data(young))
adonis(young_bray ~ Host_Taxa, data = young_df)
## Beta dispersion
beta4 <- betadisper(young_bray, young_df$Host_Taxa)
permutest(beta4, pairwise = TRUE)




################ SECTION 2 - Diversity analyses with PERMANOVA graph plotting
###################
library(devtools)
library(adespatial)
library(microbiomeSeq) #Install 'WGCNA' manually from computer if regular
installation doesn't work

### Species-specific ontogeny
Rubida <- clean_data %>% subset_samples(Host_Taxa == "Trubida")
r1 <-plot_anova_diversity(Rubida, method = c("richness", "shannon"),
                          grouping_column = "Instar", pValueCutoff = 0.05)
print(r1)

protracta <- clean_data %>% subset_samples(Host_Taxa == "Tprotracta")
r2 <- plot_anova_diversity(protracta, method = c("richness", "shannon"),
                           grouping_column = "Instar", pValueCutoff = 0.05)
print(r2)

gerst <- clean_data %>%  subset_samples(Host_Taxa == "Tgerstaeckeri")
r3 <- plot_anova_diversity(gerst, method = c("richness", "shannon"),
                           grouping_column = "Instar", pValueCutoff = 0.05)
print(r3)

lect <- clean_data %>% subset_samples(Host_Taxa == "Tlecticularia")
r4 <- plot_anova_diversity(lect, method = c("richness", "shannon"),
                           grouping_column = "Instar", pValueCutoff = 0.05)
print(r4)

sang <- clean_data %>% subset_samples(Host_Taxa == "Tsanguisuga")
r5 <- plot_anova_diversity(sang, method = c("richness", "shannon"),
                           grouping_column = "Instar", pValueCutoff = 0.1)
print(r5)



########## SECTION 3 - Full visualisation of bacterial taxa #############
#Choose level of taxa
clean_data_gen<-taxa_level(clean_data, "Genus")
#normalize the data
clean_data_barplotgen <- normalise_data(clean_data_gen, norm.method =
"relative")
```

This document was created by an application that isn't licensed to use novaPDF.
Purchase a license to generate PDF files without this notice.

```
#Different subsets:
rubida_ontogeny <- clean_data_barplotgen %>% subset_samples(Host_Taxa ==
"Trubida")
gerst_ontogeny <-  clean_data_barplotgen %>% subset_samples(Host_Taxa ==
"Tgerstaeckeri")
protra_ontogeny <- clean_data_barplotgen %>% subset_samples(Host_Taxa ==
"Tprotracta")
sang <- clean_data_barplotgen %>% subset_samples(Host_Taxa == "Tsanguisuga")
lec <- clean_data_barplotgen %>% subset_samples(Host_Taxa == "Tlecticularia")
nest2 <- clean_data_barplotgen %>% subset_samples(Locality == "Chaparral")

# Rubida only - across instars - genus level
rubida_gen <- plot_taxa(rubida_ontogeny, grouping_column = "Instar",
                        method = "hellinger", number.taxa = 20, filename =
NULL)
print(rubida_gen)

# Gerstaeckeri only - across instars - genus level
gerst_gen <- plot_taxa(gerst_ontogeny, grouping_column = "Instar",
                       method = "hellinger", number.taxa = 20, filename =
NULL)
print(gerst_gen)

## Protracta ontogeny - genus level
protra_gen <- plot_taxa(protra_ontogeny, grouping_column = "Instar",
                        method = "hellinger", number.taxa = 20, filename =
NULL)
print(protra_gen)

## Sang
sang_gen <- plot_taxa(sang, grouping_column = "Instar",
                      method = "hellinger", number.taxa = 20, filename = NULL)
print(sang_gen)

lectic_gen <- plot_taxa(lec, grouping_column = "Instar",
                        method = "hellinger", number.taxa = 20, filename =
NULL)
print(lectic_gen)

# For chaparral nest 2 - species difference - genus level
nest2gen <- plot_taxa(nest2, grouping_column = "Host_Taxa",
                      method = "hellinger", number.taxa = 20, filename = NULL)
print(nest2gen)
```

This document was created by an application that isn’t licensed to use novaPDF.
Purchase a license to generate PDF files without this notice.

```
#script for the basic data #prdouce Alpha diversity, coordination analyses and
taxonomy profile figures in supplementary files


```{r, message=FALSE}
#################### 1.0 INTRODUCTION ######################
### Get BiocManager working, and relevant packages installed
if (!requireNamespace("BiocManager", quietly = TRUE))
  install.packages("BiocManager")
BiocManager::install(c("AnnotationDbi", "DESeq2", "GO.db", "impute",
"phyloseq",
                       "preprocessCore", "biomformat", "decontam",
"microbiome"))

library(biomformat) #needed for reading biom
library(phyloseq) #needed for proper microbiome analysis
library(vegan) #community ecology analyses
library(ggplot2) #graphs
library(dplyr) #to filter and clean up the data
library(cowplot) #multiple ggplots
library(GUniFrac) #for unifrac distance based ordination
library(phangorn) #for unifrac distance based ordination using phylogenetic
tree
library(viridis) #colour package

#Read the OTU file
abund_table<-
read.csv("otu_norarefaction_abund.csv",row.names=1,check.names=FALSE)
#transpose the OTU abundance data to have sample names on rows
abund_table<-t(abund_table)
#Read in the complete meta data file
meta_table<-
read.csv("MapAndReadsUpdated2019.csv",row.names=1,check.names=FALSE)
#Read the taxonomy
OTU_taxonomy<-
read.csv("otu_norarefaction_taxa.csv",row.names=1,check.names=FALSE)



#####  1.1 - Phyloseq adaptation + cleanup
#Convert the data to phyloseq format
OTU = otu_table(as.matrix(abund_table), taxa_are_rows = FALSE)
TAX = tax_table(as.matrix(OTU_taxonomy))
SAM = sample_data(meta_table)
physeq<-merge_phyloseq(phyloseq(OTU, TAX),SAM)

clean_data<-physeq
#column names of taxonomy file
colnames(tax_table(clean_data))

#Subset field samples:
### Just nest samples
clean_nest <- clean_data %>% subset_samples(!is.na(Host_Taxa) &
                                              !is.na(T_cruzi) &
```

This document was created by an application that isn't licensed to use novaPDF.
Purchase a license to generate PDF files without this notice.
```

```
                                                              Organism == "Triatoma" & Nest !=
"houseIO" &
                                                              Nest != "bob" & Nest != "LVH" &
Origin == "field" &
                                                              T_cruzi == "N") %>%
prune_taxa(taxa_sums(.) > 0, .)
clean_nest

# Rarefy reads to even out the depth
nest_rarefy <- rarefy_even_depth(clean_nest, sample.size = 1000)
nest_rarefy


# Create some new subsets for later use here:
rubida_rarefy <- nest_rarefy %>% subset_samples(Host_Taxa == "Trubida")
chaparral <- nest_rarefy %>% subset_samples(Locality == "Chaparral")
tucson <- rubida_rarefy %>% subset_samples(Locality == "UADS" | Locality ==
"LCNCA")
young <- nest_rarefy %>% subset_samples(Instar == "L1"|Instar == "L2"|Instar
== "L3")



############## 1.2 UNCONSTRAINED ORDINATION ##############
# Set.seed for randomisation, making results reproducible
set.seed(5)
# 1.2.1 NMDS on rubida ontogeny
rubida_nmds <- ordinate(
  physeq = rubida_rarefy,
  method = "NMDS",
  distance = "bray"
)
## Stress = 0.15
## Plot it
rub2<-plot_ordination(
  physeq = rubida_rarefy,
  ordination = rubida_nmds,
  color = "Instar",
  shape = "Instar",
  title = "NMDS of Rubida ontogeny") +
  scale_color_viridis(option="plasma", discrete = TRUE, begin = 0.2, end =
0.9, direction = -1) +
  theme(panel.grid.major = element_line(size = 0.05, linetype = 'solid',
colour = "white"),
        panel.grid.minor = element_line(size = 0.05, linetype = 'solid',
colour = "white"),
        plot.background=element_rect(fill = "grey90"),
        panel.background = element_rect(fill = 'black'),
        axis.line = element_line(colour = "white"),) +
  geom_point(alpha = 1, size = 4) +
  stat_ellipse(geom='polygon', aes(colour=Instar, fill=Instar), alpha = 0.1)
print(rub2)
## 1.2.2 PERMANOVA ON ONTOGENY
rubida_bray <- phyloseq::distance(rubida_rarefy, method = "bray")
```

This document was created by an application that isn't licensed to use novaPDF.
Purchase a license to generate PDF files without this notice.

```
rubida_rarefy_df <- data.frame(sample_data(rubida_rarefy))
adonis(rubida_bray ~ Instar, data = rubida_rarefy_df, permutations = 999,
strata = rubida_rarefy_df$Locality)
beta1 <- betadisper(rubida_bray, rubida_rarefy_df$Instar, type = "median",
bias.adjust = TRUE)
permutest(beta1, pairwise = TRUE, permutations = 999, model = "direct")



####### 1.2.3 NMDS on multi-species nest from Chaparral, TX
## Now NMDS
chap_nmds <- ordinate(
  physeq = chaparral,
  method = "NMDS",
  distance = "bray"
)
## 0.09 stress
chap2<-plot_ordination(
  physeq = chaparral,
  ordination = chap_nmds,
  color = "Host_Taxa",
  shape = "Host_Taxa",
  title = "NMDS of Chaparral Multi-Species Nest") +
  scale_color_viridis(option="plasma", discrete = TRUE, begin = 0.5, end =
0.8, direction = 1) +
  theme(panel.grid.major = element_line(size = 0.02, linetype = 'solid',
colour = "black"),
        panel.grid.minor = element_line(size = 0.02, linetype = 'solid',
colour = "black"),
        plot.background=element_rect(fill = "white"),
        panel.background = element_rect(fill = 'white'),
        axis.line = element_line(colour = "black"),) +
  geom_point(alpha = 1, size = 4) +
  geom_text(aes(label=Instar),hjust=1, vjust=0, size=3, colour = "black")+
  stat_ellipse(geom='polygon', aes(colour=Host_Taxa, fill=Host_Taxa), alpha =
0.1, level = 0.95)
print(chap2)
## 1.2.4 PERMANOVA ON CHAPARRAL
chap_bray <- phyloseq::distance(chaparral, method = "bray")
chap_df <- data.frame(sample_data(chaparral))
adonis(chap_bray ~ Host_Taxa, data = chap_df, permutations = 999, strata =
chap_df$Instar) #27% of the variation is species level
## Beta dispersion
beta2 <- betadisper(chap_bray, chap_df$Host_Taxa)
permutest(beta2, pairwise = TRUE)



## 1.2.5 NMDS on rubida from 2 locations in Arizona
tuc_nmds <- ordinate(
  physeq = tucson,
  method = "NMDS",
  distance = "bray"
)
```

This document was created by an application that isn't licensed to use novaPDF.
Purchase a license to generate PDF files without this notice.

```
## 0.17 stress
tuc1<-plot_ordination(
  physeq = tucson,
  ordination = tuc_nmds,
  color = "Locality",
  shape = "Instar_range",
  title = "NMDS of Tucson rubida") +
  scale_color_viridis(option = "plasma", discrete = TRUE, begin = 0.3, end =
0.8)+
  theme(panel.grid.major = element_line(size = 0.02, linetype = 'solid',
colour = "black"),
        panel.grid.minor = element_line(size = 0.02, linetype = 'solid',
colour = "black"),
        plot.background=element_rect(fill = "gray90"),
        panel.background = element_rect(fill = 'white'),
        axis.line = element_line(colour = "black"),) +
  geom_point(alpha = 1, size = 4) +
  #geom_text(aes(label=Instar),hjust=1, vjust=0, size=3, colour = "black")+
  stat_ellipse(geom='polygon', aes(colour=Locality, fill=Locality), alpha =
0.1, level = 0.95)
print(tuc1)
## 1.2.6. PERMANOVA ON TUCSON NEST RUBIDA
tuc_bray <- phyloseq::distance(tucson, method = "bray")
tuc_df <- data.frame(sample_data(tucson))
adonis(tuc_bray ~ Locality*Instar, data = tuc_df)
beta3 <- betadisper(tuc_bray, tuc_df$Locality)
permutest(beta3)



######  1.2.7 NMDS on young instar species-specific differences
young_nmds <- ordinate(
  physeq = young,
  method = "NMDS",
  distance = "bray"
)
### stress = ~0.2
young1<-plot_ordination(
  physeq = young,
  ordination = young_nmds,
  color = "Host_Taxa",
  shape = "Host_Taxa",
  title = "Young instars") +
  scale_color_viridis(option = "plasma", discrete = TRUE, begin = 0.2, end =
0.9)+
  theme(panel.grid.major = element_line(size = 0.02, linetype = 'solid',
colour = "black"),
        panel.grid.minor = element_line(size = 0.02, linetype = 'solid',
colour = "black"),
        plot.background=element_rect(fill = "white"),
        panel.background = element_rect(fill = 'white'),
        axis.line = element_line(colour = "black"),) +
  geom_point(alpha = 1, size = 4)+
```

This document was created by an application that isn’t licensed to use novaPDF.
Purchase a license to generate PDF files without this notice.

```
  stat_ellipse(geom='polygon', aes(colour=Host_Taxa, fill=Host_Taxa), alpha =
0.1, level = 0.95)
print(young1)
## 1.2.8
young_bray <- phyloseq::distance(young, method = "bray")
young_df <- data.frame(sample_data(young))
adonis(young_bray ~ Host_Taxa, data = young_df)
## Beta dispersion
beta4 <- betadisper(young_bray, young_df$Host_Taxa)
permutest(beta4, pairwise = TRUE)




################ SECTION 2 - Diversity analyses with PERMANOVA graph plotting
###################
library(devtools)
library(adespatial)
library(microbiomeSeq) #Install 'WGCNA' manually from computer if regular
installation doesn't work

### Species-specific ontogeny
Rubida <- clean_data %>% subset_samples(Host_Taxa == "Trubida")
r1 <-plot_anova_diversity(Rubida, method = c("richness", "shannon"),
                          grouping_column = "Instar", pValueCutoff = 0.05)
print(r1)

protracta <- clean_data %>% subset_samples(Host_Taxa == "Tprotracta")
r2 <- plot_anova_diversity(protracta, method = c("richness", "shannon"),
                           grouping_column = "Instar", pValueCutoff = 0.05)
print(r2)

gerst <- clean_data %>%  subset_samples(Host_Taxa == "Tgerstaeckeri")
r3 <- plot_anova_diversity(gerst, method = c("richness", "shannon"),
                           grouping_column = "Instar", pValueCutoff = 0.05)
print(r3)

lect <- clean_data %>% subset_samples(Host_Taxa == "Tlecticularia")
r4 <- plot_anova_diversity(lect, method = c("richness", "shannon"),
                           grouping_column = "Instar", pValueCutoff = 0.05)
print(r4)

sang <- clean_data %>% subset_samples(Host_Taxa == "Tsanguisuga")
r5 <- plot_anova_diversity(sang, method = c("richness", "shannon"),
                           grouping_column = "Instar", pValueCutoff = 0.1)
print(r5)



########## SECTION 3 - Full visualisation of bacterial taxa ############
#Choose level of taxa
clean_data_gen<-taxa_level(clean_data, "Genus")
#normalize the data
```

This document was created by an application that isn’t licensed to use novaPDF.
Purchase a license to generate PDF files without this notice.

```
clean_data_barplotgen <- normalise_data(clean_data_gen, norm.method =
"relative")

#Different subsets:
rubida_ontogeny <- clean_data_barplotgen %>% subset_samples(Host_Taxa ==
"Trubida")
gerst_ontogeny <-  clean_data_barplotgen %>% subset_samples(Host_Taxa ==
"Tgerstaeckeri")
protra_ontogeny <- clean_data_barplotgen %>% subset_samples(Host_Taxa ==
"Tprotracta")
sang <- clean_data_barplotgen %>% subset_samples(Host_Taxa == "Tsanguisuga")
lec <- clean_data_barplotgen %>% subset_samples(Host_Taxa == "Tlecticularia")
nest2 <- clean_data_barplotgen %>% subset_samples(Locality == "Chaparral")

# Rubida only - across instars - genus level
rubida_gen <- plot_taxa(rubida_ontogeny, grouping_column = "Instar",
                        method = "hellinger", number.taxa = 20, filename =
NULL)
print(rubida_gen)

# Gerstaeckeri only - across instars - genus level
gerst_gen <- plot_taxa(gerst_ontogeny, grouping_column = "Instar",
                       method = "hellinger", number.taxa = 20, filename =
NULL)
print(gerst_gen)

## Protracta ontogeny - genus level
protra_gen <- plot_taxa(protra_ontogeny, grouping_column = "Instar",
                        method = "hellinger", number.taxa = 20, filename =
NULL)
print(protra_gen)

## Sang
sang_gen <- plot_taxa(sang, grouping_column = "Instar",
                      method = "hellinger", number.taxa = 20, filename = NULL)
print(sang_gen)

lectic_gen <- plot_taxa(lec, grouping_column = "Instar",
                        method = "hellinger", number.taxa = 20, filename =
NULL)
print(lectic_gen)

# For chaparral nest 2 - species difference - genus level
nest2gen <- plot_taxa(nest2, grouping_column = "Host_Taxa",
                      method = "hellinger", number.taxa = 20, filename = NULL)
print(nest2gen)
```

```
```
```

This document was created by an application that isn’t licensed to use novaPDF.
Purchase a license to generate PDF files without this notice.
```

```
#script for the decontam data #prdouce Alpha diversity, coordination analyses
and taxonomy profile figures in supplementary files



```{r, message=FALSE}

#################### 1.0 INTRODUCTION #####################
rm(list=ls())
### Get BiocManager working, and relevant packages installed
if (!requireNamespace("BiocManager", quietly = TRUE))
  install.packages("BiocManager")
BiocManager::install(c("AnnotationDbi", "DESeq2", "GO.db", "impute",
"phyloseq",
                       "preprocessCore", "biomformat", "decontam",
"microbiome"))

library(biomformat) #needed for reading biom
library(phyloseq) #needed for proper microbiome analysis
library(vegan) #community ecology analyses
library(ggplot2) #graphs
library(dplyr) #to filter and clean up the data
library(cowplot) #multiple ggplots
library(GUniFrac) #for unifrac distance based ordination
library(phangorn) #for unifrac distance based ordination using phylogenetic
tree
library(viridis) #colour package

#Read the OTU file
abund_table<-read.csv("otu_decontam_abund.csv",row.names=1,check.names=FALSE)
#transpose the OTU abundance data to have sample names on rows
abund_table<-t(abund_table)
#Read in the complete meta data file
meta_table<-
read.csv("MapAndReadsUpdated2019.csv",row.names=1,check.names=FALSE)
#Read the taxonomy
OTU_taxonomy<-read.csv("otu_decontam_taxa.csv",row.names=1,check.names=FALSE)


colnames(meta_table)<-c("ReadCount", "Plate", "WellPosition", "Organism",
"State",
                        "Tcruzi","Host_Taxa", "Instar", "Instar_range", "Sex",
"Locality",
                        "Nest", "Latitude", "Longitude", "Sample")


#####  1.1 - Phyloseq adaptation + cleanup
#Convert the data to phyloseq format
OTU = otu_table(as.matrix(abund_table), taxa_are_rows = FALSE)
TAX = tax_table(as.matrix(OTU_taxonomy))
SAM = sample_data(meta_table)
physeq<-merge_phyloseq(phyloseq(OTU, TAX),SAM,OTU_tree,OTU_seqs)
```

This document was created by an application that isn't licensed to use novaPDF.
Purchase a license to generate PDF files without this notice.
```

```
clean_data<-physeq
#column names of taxonomy file
colnames(tax_table(clean_data))


#Subset field samples:
# Just nest samples
clean_nest <- clean_data %>% subset_samples(!is.na(Host_Taxa) &
                                              !is.na(T_cruzi) &
                                              Organism == "Triatoma" & Nest !=
"houseIO" &
                                              Nest != "bob" & Nest != "LVH" &
Origin == "field" &
                                              T_cruzi == "N") %>%
prune_taxa(taxa_sums(.) > 0, .)
clean_nest

###### Removing low abundance OTUs
# melt to long format (for ggploting)
# prune out low abundance genera
nest_genus <- clean_nest %>%
  tax_glom(taxrank = "Genus") %>%                      # agglomerate at genus
level
  transform_sample_counts(function(x) {x/sum(x)} ) %>% # Transform to rel.
abundance
  psmelt() %>%                                         # Melt to long format
  filter(Abundance > 0.01) %>%                          # Filter out low
abundance taxa
  arrange(Sample)                                      # Arrange
alphabetically by sample

# Rarefy reads to even out the depth
nest_rarefy <- rarefy_even_depth(clean_nest, sample.size = 1000)
nest_rarefy

# Create some new subsets for later use here:
rubida_rarefy <- nest_rarefy %>% subset_samples(Host_Taxa == "Trubida")
chaparral <- nest_rarefy %>% subset_samples(Locality == "Chaparral")
tucson <- rubida_rarefy %>% subset_samples(Locality == "UADS" | Locality ==
"LCNCA")
young <- nest_rarefy %>% subset_samples(Instar == "L1"|Instar == "L2"|Instar
== "L3")

############## 1.2 UNCONSTRAINED ORDINATION ##############
# Set.seed for randomisation, making results reproducible
set.seed(5)
# 1.2.1 NMDS on rubida ontogeny
rubida_nmds <- ordinate(
  physeq = rubida_rarefy,
  method = "NMDS",
  distance = "bray"
)
## Stress = 0.15
## Plot it
```

This document was created by an application that isn't licensed to use novaPDF.
Purchase a license to generate PDF files without this notice.

```
rub2<-plot_ordination(
  physeq = rubida_rarefy,
  ordination = rubida_nmds,
  color = "Instar",
  shape = "Instar",
  title = "NMDS of Rubida ontogeny") +
  scale_color_viridis(option="plasma", discrete = TRUE, begin = 0.2, end =
0.9, direction = -1) +
  theme(panel.grid.major = element_line(size = 0.05, linetype = 'solid',
colour = "white"),
        panel.grid.minor = element_line(size = 0.05, linetype = 'solid',
colour = "white"),
        plot.background=element_rect(fill = "grey90"),
        panel.background = element_rect(fill = 'black'),
        axis.line = element_line(colour = "white"),) +
  geom_point(alpha = 1, size = 4) +
  stat_ellipse(geom='polygon', aes(colour=Instar, fill=Instar), alpha = 0.1)
print(rub2)
## 1.2.2 PERMANOVA ON ONTOGENY
rubida_bray <- phyloseq::distance(rubida_rarefy, method = "bray")
rubida_rarefy_df <- data.frame(sample_data(rubida_rarefy))
adonis(rubida_bray ~ Instar, data = rubida_rarefy_df, permutations = 999,
strata = rubida_rarefy_df$Locality)
beta1 <- betadisper(rubida_bray, rubida_rarefy_df$Instar, type = "median",
bias.adjust = TRUE)
permutest(beta1, pairwise = TRUE, permutations = 999, model = "direct")



####### 1.2.3 NMDS on multi-species nest from Chaparral, TX
## Now NMDS
chap_nmds <- ordinate(
  physeq = chaparral,
  method = "NMDS",
  distance = "bray"
)
## 0.09 stress
chap2<-plot_ordination(
  physeq = chaparral,
  ordination = chap_nmds,
  color = "Host_Taxa",
  shape = "Host_Taxa",
  title = "NMDS of Chaparral Multi-Species Nest") +
  scale_color_viridis(option="plasma", discrete = TRUE, begin = 0.5, end =
0.8, direction = 1) +
  theme(panel.grid.major = element_line(size = 0.02, linetype = 'solid',
colour = "black"),
        panel.grid.minor = element_line(size = 0.02, linetype = 'solid',
colour = "black"),
        plot.background=element_rect(fill = "white"),
        panel.background = element_rect(fill = 'white'),
        axis.line = element_line(colour = "black"),) +
  geom_point(alpha = 1, size = 4) +
  geom_text(aes(label=Instar),hjust=1, vjust=0, size=3, colour = "black")+
```

This document was created by an application that isn’t licensed to use novaPDF.
Purchase a license to generate PDF files without this notice.

```
  stat_ellipse(geom='polygon', aes(colour=Host_Taxa, fill=Host_Taxa), alpha =
0.1, level = 0.95)
print(chap2)
## 1.2.4 PERMANOVA ON CHAPARRAL
chap_bray <- phyloseq::distance(chaparral, method = "bray")
chap_df <- data.frame(sample_data(chaparral))
adonis(chap_bray ~ Host_Taxa, data = chap_df, permutations = 999, strata =
chap_df$Instar) #27% of the variation is species level
## Beta dispersion
beta2 <- betadisper(chap_bray, chap_df$Host_Taxa)
permutest(beta2, pairwise = TRUE)



## 1.2.5 NMDS on rubida from 2 locations in Arizona
tuc_nmds <- ordinate(
  physeq = tucson,
  method = "NMDS",
  distance = "bray"
)
## 0.17 stress
tuc1<-plot_ordination(
  physeq = tucson,
  ordination = tuc_nmds,
  color = "Locality",
  shape = "Instar_range",
  title = "NMDS of Tucson rubida") +
  scale_color_viridis(option = "plasma", discrete = TRUE, begin = 0.3, end =
0.8)+
  theme(panel.grid.major = element_line(size = 0.02, linetype = 'solid',
colour = "black"),
        panel.grid.minor = element_line(size = 0.02, linetype = 'solid',
colour = "black"),
        plot.background=element_rect(fill = "gray90"),
        panel.background = element_rect(fill = 'white'),
        axis.line = element_line(colour = "black"),) +
  geom_point(alpha = 1, size = 4) +
  #geom_text(aes(label=Instar),hjust=1, vjust=0, size=3, colour = "black")+
  stat_ellipse(geom='polygon', aes(colour=Locality, fill=Locality), alpha =
0.1, level = 0.95)
print(tuc1)
## 1.2.6. PERMANOVA ON TUCSON NEST RUBIDA
tuc_bray <- phyloseq::distance(tucson, method = "bray")
tuc_df <- data.frame(sample_data(tucson))
adonis(tuc_bray ~ Locality*Instar, data = tuc_df)
beta3 <- betadisper(tuc_bray, tuc_df$Locality)
permutest(beta3)



######  1.2.7 NMDS on young instar species-specific differences
young_nmds <- ordinate(
  physeq = young,
  method = "NMDS",
```

This document was created by an application that isn't licensed to use novaPDF.
Purchase a license to generate PDF files without this notice.

```
  distance = "bray"
)
### stress = ~0.2
young1<-plot_ordination(
  physeq = young,
  ordination = young_nmds,
  color = "Host_Taxa",
  shape = "Host_Taxa",
  title = "Young instars") +
  scale_color_viridis(option = "plasma", discrete = TRUE, begin = 0.2, end =
0.9)+
  theme(panel.grid.major = element_line(size = 0.02, linetype = 'solid',
colour = "black"),
        panel.grid.minor = element_line(size = 0.02, linetype = 'solid',
colour = "black"),
        plot.background=element_rect(fill = "white"),
        panel.background = element_rect(fill = 'white'),
        axis.line = element_line(colour = "black"),) +
  geom_point(alpha = 1, size = 4)+
  stat_ellipse(geom='polygon', aes(colour=Host_Taxa, fill=Host_Taxa), alpha =
0.1, level = 0.95)
print(young1)
## 1.2.8
young_bray <- phyloseq::distance(young, method = "bray")
young_df <- data.frame(sample_data(young))
adonis(young_bray ~ Host_Taxa, data = young_df)
## Beta dispersion
beta4 <- betadisper(young_bray, young_df$Host_Taxa)
permutest(beta4, pairwise = TRUE)



################ SECTION 2 - Diversity analyses with PERMANOVA graph plotting
###################
library(devtools)
library(adespatial)
library(microbiomeSeq) #Install 'WGCNA' manually from computer if regular
installation doesn't work

### Species-specific ontogeny
Rubida <- clean_data %>% subset_samples(Host_Taxa == "Trubida")
r1 <-plot_anova_diversity(Rubida, method = c("richness", "shannon"),
                          grouping_column = "Instar", pValueCutoff = 0.05)
print(r1)

protracta <- clean_data %>% subset_samples(Host_Taxa == "Tprotracta")
r2 <- plot_anova_diversity(protracta, method = c("richness", "shannon"),
                           grouping_column = "Instar", pValueCutoff = 0.05)
print(r2)

gerst <- clean_data %>%  subset_samples(Host_Taxa == "Tgerstaeckeri")
r3 <- plot_anova_diversity(gerst, method = c("richness", "shannon"),
                           grouping_column = "Instar", pValueCutoff = 0.05)
```

This document was created by an application that isn’t licensed to use novaPDF.
Purchase a license to generate PDF files without this notice.

```
print(r3)

lect <- clean_data %>% subset_samples(Host_Taxa == "Tlecticularia")
r4 <- plot_anova_diversity(lect, method = c("richness", "shannon"),
                           grouping_column = "Instar", pValueCutoff = 0.05)
print(r4)

sang <- clean_data %>% subset_samples(Host_Taxa == "Tsanguisuga")
r5 <- plot_anova_diversity(sang, method = c("richness", "shannon"),
                           grouping_column = "Instar", pValueCutoff = 0.1)
print(r5)



########## SECTION 3 - Full visualisation of bacterial taxa ############
#Choose level of taxa
clean_data_gen<-taxa_level(clean_data, "Genus")
#normalize the data
clean_data_barplotgen <- normalise_data(clean_data_gen, norm.method =
"relative")

#Different subsets:
rubida_ontogeny <- clean_data_barplotgen %>% subset_samples(Host_Taxa ==
"Trubida")
gerst_ontogeny <-  clean_data_barplotgen %>% subset_samples(Host_Taxa ==
"Tgerstaeckeri")
protra_ontogeny <- clean_data_barplotgen %>% subset_samples(Host_Taxa ==
"Tprotracta")
sang <- clean_data_barplotgen %>% subset_samples(Host_Taxa == "Tsanguisuga")
lec <- clean_data_barplotgen %>% subset_samples(Host_Taxa == "Tlecticularia")
nest2 <- clean_data_barplotgen %>% subset_samples(Locality == "Chaparral")

# Rubida only - across instars - genus level
rubida_gen <- plot_taxa(rubida_ontogeny, grouping_column = "Instar",
                        method = "hellinger", number.taxa = 20, filename =
NULL)
print(rubida_gen)

# Gerstaeckeri only - across instars - genus level
gerst_gen <- plot_taxa(gerst_ontogeny, grouping_column = "Instar",
                       method = "hellinger", number.taxa = 20, filename =
NULL)
print(gerst_gen)

## Protracta ontogeny - genus level
protra_gen <- plot_taxa(protra_ontogeny, grouping_column = "Instar",
                        method = "hellinger", number.taxa = 20, filename =
NULL)
print(protra_gen)

## Sang
sang_gen <- plot_taxa(sang, grouping_column = "Instar",
                      method = "hellinger", number.taxa = 20, filename = NULL)
print(sang_gen)
```

This document was created by an application that isn't licensed to use novaPDF.
Purchase a license to generate PDF files without this notice.

```
lectic_gen <- plot_taxa(lec, grouping_column = "Instar",
                        method = "hellinger", number.taxa = 20, filename =
NULL)
print(lectic_gen)

# For chaparral nest 2 - species difference - genus level
nest2gen <- plot_taxa(nest2, grouping_column = "Host_Taxa",
                      method = "hellinger", number.taxa = 20, filename = NULL)
print(nest2gen)
```

#script for mantel test in main text

```{r, message=FALSE}

library(biomformat)
library(vegan)
library(ecodist)

ototab <- "UltracleanAllSamples_500.biom"

#ADD WHERE NEEDED THE SCRIPT FOR SAMPLE FILTERING
otutab <- otutab %>% subset_samples(Host_Taxa == "Trubida", Instar_range ==
"L1-3", Locality == "UADS")

distmat <- "distance_matrix_qgis.csv"

dist <- read.table(distmat, header=TRUE, sep = ",", row.names = 1, as.is=TRUE)
distm <- as.matrix(dist)

otu.biom <- read_biom(otutab)
otu.table <- t(as.data.frame(as(biom_data(otu.biom), "matrix")))
dissi <- vegdist(otu_table(otu.table), method = "bray")
dissim <- as.matrix(dissi)

mantel(as.dist(dissim) ~ as.dist(distm), mrank = TRUE)

```

This document was created by an application that isn't licensed to use novaPDF.
Purchase a license to generate PDF files without this notice.

### evaluation of phylogenetic constraint on the microbiomes

````
```{python, message=FALSE}

#using QIIME 1 scripts
#map94.txt is 94 L1_3 samples with available coxB sequence: https://
www.dropbox.com/s/hov5c3ljr16b9rd/map94.txt?dl=0

 filter_samples_from_otu_table.py -i ../Ultraclean500MapNovember.biom -o
L1_3_set94.biom -m map94.txt -s 'Instar:*'

#calculate Bray Curtis distances for microbiome data

 beta_diversity.py -i L1_3_set94.biom -m bray_curtis -o ./beta_div_set94

#calculate phylogenetic distances between Triatoma individuals using NJ
algorithm in Geneious

#run Mantel test for assessing correlation between phylogenetic and microbiome
distances, inputs: https://www.dropbox.com/s/5fkpq5irhend1if/tn94f.txt?dl=0
AND https://www.dropbox.com/s/uplzi7gkg3uicgq/bc94f.txt?dl=0

 compare_distance_matrices.py --method mantel -i tn94f.txt,bc94f.txt -o
mantel_out_set94 -n 999




```
````

#script for checking homegenity condition between two instar ranges
#betadispersion table and bar plots are presented in supplementary files

````
```{r, message=FALSE}
#load libraries
library(phyloseq); packageVersion("phyloseq")
library(ggplot2); packageVersion("ggplot2")
library(dplyr); packageVersion("dplyr") #require to filter and clean up the
data
library(vegan); packageVersion("vegan")


#upload the ultraclean files

#upload the ultraclean OTU abundance file
````

This document was created by an application that isn't licensed to use novaPDF.
Purchase a license to generate PDF files without this notice.
````

```
ultra_abund_table<-
read.csv("otu_ultraclean_abund.csv",row.names=1,check.names=FALSE)

#transpose the OTU abundance data to have sample names on rows
ultra_abund_table<-t(ultra_abund_table)

#upload the meta data file
ultra_meta_table<-
read.csv("ultrameta_28022020.csv",row.names=1,check.names=FALSE)


#load the taxonomy
ultra_OTU_taxonomy<-
read.csv("otu_ultraclean_taxa.csv",row.names=1,check.names=FALSE)

#Convert the data to phyloseq format
ultra_OTU = otu_table(as.matrix(ultra_abund_table), taxa_are_rows = FALSE)
ultra_TAX = tax_table(as.matrix(ultra_OTU_taxonomy))
ultra_SAM = sample_data(ultra_meta_table)
ultra_physeq<-merge_phyloseq(phyloseq(ultra_OTU, ultra_TAX),ultra_SAM)
ultra_physeq

#remove "NA" in Hosttaxa_instarrange of ultra_physeq
ultra_physeq <- ultra_physeq %>%
  subset_samples(Host_instarrange != "NA") %>%
  prune_taxa(taxa_sums(.) > 0, .)

#if need to be renamed #here i renamed Domain into Kingdom
colnames(tax_table(ultra_physeq)) <- c("Kingdom", "Phylum", "Class", "Order",
"Family", "Genus", "Species")

# Scale reads to even depth
ultra_physeq_rarefy <- rarefy_even_depth(ultra_physeq, sample.size = 1000)

ultra_physeq_rarefy



#ordination analyses using jaccard

ultra_physeq_rarefy_jaccard <- distance(ultra_physeq_rarefy, "jaccard")
ultra_physeq_rarefy_df <- as(sample_data(ultra_physeq_rarefy), "data.frame")
p2 <- plot_ordination(ultra_physeq_rarefy, ultra_physeq_rarefy_jaccard, color
= "Instar_range")
p2 + theme_bw() + theme(text = element_text(size = 16)) + geom_point(size = 4)
+ stat_ellipse(aes(group =Instar_range))
p2


#Permanova for the instar range in whole ultra dataset

adonis_Instar_range <- adonis(ultra_physeq_rarefy_jaccard ~ Instar_range, data
= ultra_physeq_rarefy_df)
adonis_Instar_range
```

This document was created by an application that isn’t licensed to use novaPDF.
Purchase a license to generate PDF files without this notice.

````
#betadiseprsion for the instar range in whole dataset
groups <- ultra_physeq_rarefy_df[["Instar_range"]]
mod <- betadisper(ultra_physeq_rarefy_jaccard, groups)
anova(mod)

#the dispersion is different between groups, then examine
plot(mod)
boxplot(mod)
mod.HSD <- TukeyHSD(mod )
plot(mod.HSD)


```
````

This document was created by an application that isn’t licensed to use novaPDF.
Purchase a license to generate PDF files without this notice.
````
